# Rapid Histone Post-Translational Modification Analysis Using Alternative Proteases and Tandem Mass Tags

**DOI:** 10.64898/2026.02.13.705817

**Authors:** Natalie P. Turner, Sabyasachi Baboo, Patrick Garrett, Jolene K. Diedrich, Michal Bajo, Marisa Roberto, John R. Yates

## Abstract

Histone post-translational modifications (PTMs) alter chromatin dynamics and contribute to the regulation of gene expression in health and disease, yet mass spectrometry-based histone PTM analysis remains constrained by inefficient sample preparation workflows. Here, we develop RIPUP (*R*apid *I*dentification of histone *P*TMs in *U*nderivatized *P*eptides), a streamlined multi-protease workflow that reduces sample preparation to hours while improving PTM coverage and quantitative accuracy. Systematic evaluation of Arg-C Ultra and a recombinant (r)-Chymotrypsin protease under varied conditions, including standard derivatization with propionic anhydride and tandem mass tag (TMT) labeling, demonstrated that Arg-C Ultra with TMT labeling achieves a detection of total PTM that exceeds Trypsin-based approaches. Using the HiP-Frag computational framework for unrestrictive PTM identification, we discovered that TMT’s tertiary amine provides charge compensation that rescues the ionization of negatively charged acylations revealing 58 succinylation and 31 glutarylation sites – a ‘dark epigenome’ largely undetected by propionylation-based methods. Complementary digestion with Arg-C Ultra and r-Chymotrypsin provides orthogonal sequence coverage, enabling detection of PTMs in H2A variants, linker histones, and regions poorly represented by arginine-specific cleavage alone. In HEK293T cells treated with the pan-sirtuin inhibitor nicotinamide, RIPUP quantified 112 statistically significant peptidoforms (adj *p* < 0.05), predominantly increasing with NAM dose (88 up, 24 down). Application of RIPUP to frozen-thawed rat hippocampal sections within a 3-hour workflow identified >200 PTMs including H3 K27/K36/K37 methylation, H4 N-terminal acetylation patterns, and H2A K118/K119 ubiquitination. This rapid, high-efficiency platform enables timely discovery of epigenetic mechanisms and accelerates the path from PTM identification to therapeutic target validation.

## Background

Mass spectrometry (MS)-based proteomics is a popular and powerful tool for analyzing histone post-translational modifications (PTMs), known as the ‘histone code’ ^1,2^. Histone proteins assemble as octamers comprised of two copies of histone H2A, H2B, H3 and H4 to form the nucleosome core, which is sequentially wrapped in ∼146-147 bp DNA during nucleosome assembly ^3^. Tightly wound nucleosomes form chromatin, the major structural unit of chromosomes. Fundamental aspects of epigenetic regulation are under the control of ‘writers’ and ‘erasers’ that add and remove chemical groups (PTMs) to histones. These changes control gene transcription by altering chromatin structure and thus the accessibility of transcriptional machinery to chromatin. Histone PTM research has focused on understanding nuanced epigenetic mechanisms implicated in health and disease paradigms, including cancer ^3^, neurobiology ^4^, addiction ^5^, and the gut microbiome ^6^.

In 2007, Garcia et al. ^2^ significantly advanced the field of MS-based histone PTM analysis by introducing a workflow that includes a chemical derivatization step to propionylate ε-amino groups of lysine (K) and peptide N-termini. This modification prevents Trypsin cleavage after K, generating longer, more hydrophobic peptides, improving solid-phase retention and chromatographic separation ^2,7^. These tryptic peptides contain R as the C-terminal amino acid and produce stable, singly charged *y*-ion series fragments. While this workflow has undergone multiple iterations since its initial publication ^7–9^, the optimization of parameters that may improve workflow efficiency, peptide sequence coverage, reproducibility, lower peptide coefficients of variation (CVs), or identify novel PTMs is still relatively under-explored ^9–12^. Importantly, the propionylation-based protocol remains one of the most influential in the field to date and has recently been applied for the analysis of histone PTMs in extracellular vesicles ^13^ and single-cells ^14,15^.

Two new-to-market enzymes – a recombinant (r)-Chymotrypsin and Arg-C Ultra (Promega™) – offer significant advantages to the standard Trypsin and propionylation approach for histone PTM analysis by MS-based proteomics ^10,16^. r-Chymotrypsin and Arg-C Ultra require shorter incubation times (2 h compared to >6 h with Trypsin) and do not require blocking of K residues as neither Arg-C Ultra nor r-Chymotrypsin cleaves at the lysine C-terminus. If derivatization is preferred for its chromatographic benefits, propionylation is required after digestion only to label unmodified lysines and peptide N-termini. In contrast to standard Chymotrypsin, r-Chymotrypsin does not cleave after tryptophan (W) and shows increased specificity for phenylalanine (F), tyrosine (Y), and leucine (L), generating different peptides to Trypsin and Arg-C Ultra digestion, thus capturing different segments of histone protein sequences. These changes to the established protocol ^1,2^ can significantly reduce the overall sample preparation time for development of high-throughput workflows and reveal insights into histone PTMs not seen with Trypsin digestion ^10^.

Other approaches for labeling histone peptides have been reported, including recent work by Ryzhaya et al. ^10^, which demonstrated that the Arg-C Ultra protease, which has substantially better arginine-cleavage specificity and efficiency than conventional Arg-C, can be combined with peptide-level derivatization using trimethylacetic anhydride (TMA) to reduce histone sample preparation time to ∼3-4 hours. Similarly, tandem mass tags (TMT)^17^ is a well-established amine-reactive label that has not been systematically evaluated for histone PTM analysis. TMT-labeling adds isobaric tags to primary amines, such as those of post-digestion peptide N-termini and unmodified K residues. Like propionyl labeling, TMT labeling creates peptides that are more hydrophobic, but with potentially higher labeling efficiency in a single one- hour incubation step ^18,19^. Additionally, TMT is compatible with multiplexed data acquisition and may offer benefits for quantitative analysis.

Vai et al. (2025) ^20^ recently developed HiP-Frag, an unrestrictive search workflow that identified 60 previously unreported modifications on core histones and 13 on linker histones across multiple cell lines and tissue samples. While these computational advances have revealed the existence of numerous uncommon modifications, the key question of whether different sample preparation and chemical labeling strategies preferentially detect certain PTM classes remain unaddressed and has implications for potentially biasing our view of the epigenetic landscape. This consideration is particularly important given that propionylation neutralizes positive charges on lysine residues, which may affect the ionization efficiency of peptides bearing negatively charged modifications.

To address these challenges, we performed a comprehensive systematic evaluation of both the standard Trypsin protocol ^1^ and two alternative proteases, Arg-C Ultra and r-Chymotrypsin (Promega™), under multiple experimental conditions using histones extracted from HEK293T cells. We tested each protease with and without propionylation, under denaturing and non-denaturing conditions, generating 10 distinct conditions (Figure 1). We also assessed TMT labeling as an alternative to derivatization with propionic anhydride. In addition to confirming recent reports of Arg-C Ultra’s superior specificity, our analysis revealed that TMT labeling provides unique advantages for detecting negatively charged acylations through charge compensation at the peptide N-terminus. We further demonstrated that r-Chymotrypsin provides complementary sequence coverage for H2A variants and linker histones that are poorly represented by arginine-specific cleavage. We performed a quantitative experiment using a pan-sirtuin inhibitor to induce global acetylation increase and found that Arg-C Ultra is particularly robust for detecting differences in multiply modified peptidoforms. As a proof-of-concept, we performed our streamlined dual-protease protocol, RIPUP (*R*apid *I*dentification of histone *P*TMs in *U*nderivatized *P*eptides), on histones extracted from rat hippocampal sections to identify peptidoforms and PTM sites of biological interest in under 3 h of sample preparation time.

**Figure 1.**
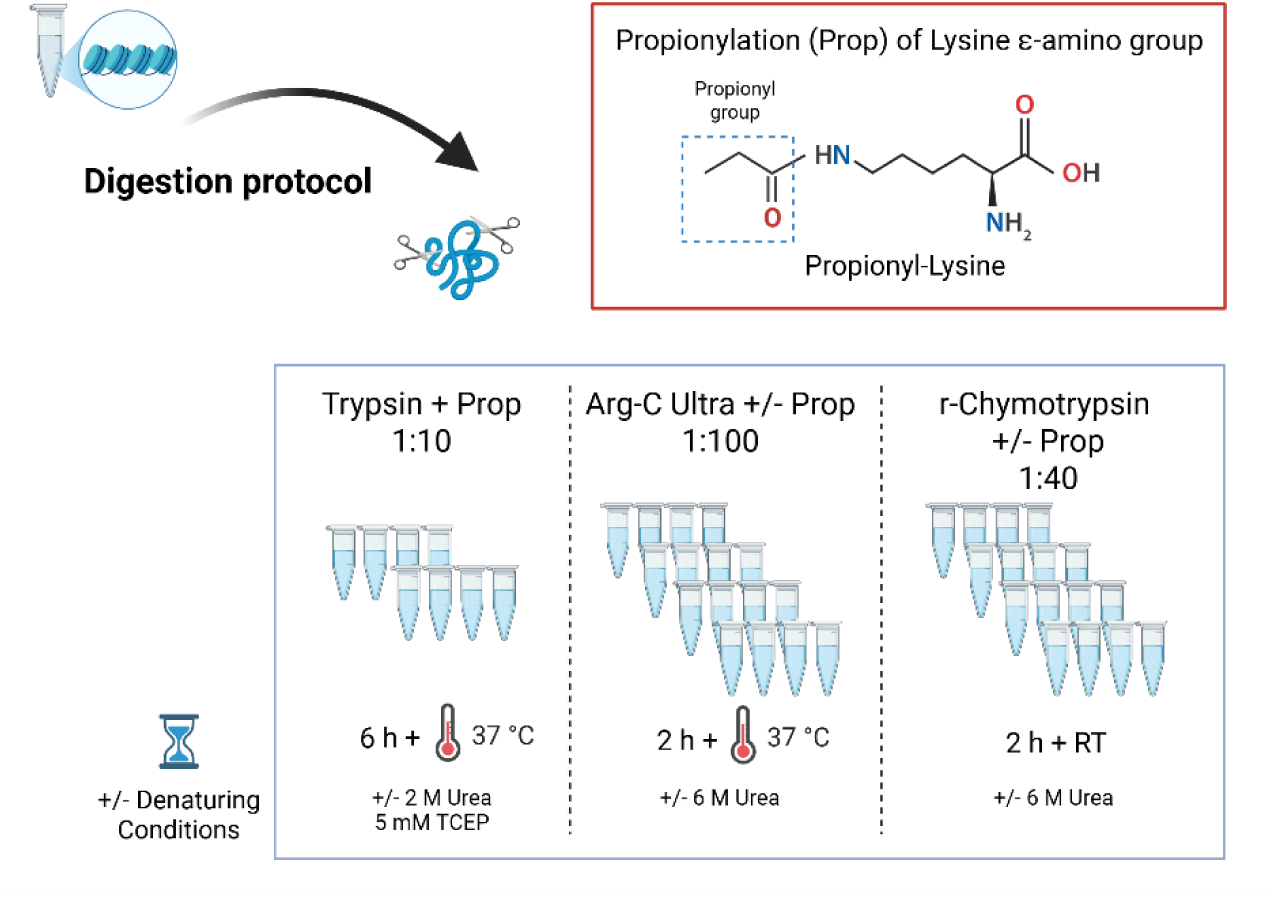
Experimental workflow for alternative protease and digestion conditions comparison. Extracted histones were subjected to enzymatic digestion using three different protease conditions with or without chemical propionylation (Prop) and denaturing agent (2 or 6 M urea). Addition of a propionyl group to lysine ε-amino groups neutralizes positive charges but prevents Trypsin cleavage at lysine residues, enabling retention of shorter peptides for higher sampling of histone PTMs. The conventional Trypsin-based workflow with propionylation at 1:10 enzyme-to-substrate ratio, requires 6 h incubation at 37 °C. Samples were prepared with or without denaturing conditions. Arg-C Ultra digestion at 1:100 enzyme-to-substrate ratio with or without propionylation, requires only 2 h at 37 °C. Recombinant chymotrypsin (r-Chymotrypsin) digestion at 1:40 enzyme-to-substrate ratio with or without propionylation, was performed at room temperature (RT) for 2 h.

## Methods

Adult male Sprague-Dawley rats used in this study (Charles River Laboratories, Raleigh, NC) were kept in accordance with the ARRIVE guidelines and were approved by the Scripps Research Institute (TSRI) Animal Care and Use Committee (IACUC #09-0006), consistent with the National Institutes of Health Guide for the Care and Use of Laboratory Animals. The animals were housed in a temperature- and humidity-controlled room (12 h reverse light cycle) and provided with food and water *ad libitum*. The rats (*n* = 5, 446 ± 17.8 g) were anesthetized with isoflurane (3%), decapitated, and the brains were rapidly removed and placed into ice-cold high-sucrose cutting solution (206.0 mM sucrose, 2.5 mM KCl, 0.5 mM CaCl_2_, 7.0 mM MgCl_2_, 1.2 mM NaH_2_PO_4_, 26 mM NaHCO_3_, 5.0 mM glucose, and 5 mM HEPES) gassed with 95% O_2_ and 5% CO_2_ ^21,22^. A Vibrotome VS1000 (Leica Microsystems) was used to cut 300 μm coronal slices containing hippocampus (AP: -2.00 to -3.25 from bregma). The slices were transferred to cold, oxygenated (95% O_2_ and 5% CO_2_) artificial cerebrospinal fluid (130 mM NaCl, 3.5 mM KCl, 1.25 mM NaH_2_PO_4_, 1.5 mM MgSO_4_, 2 mM CaCl_2_, 24 mM NaHCO_3_, and 10 mM glucose) and hippocampi were isolated and collected in 1.5 mL microcentrifuge tubes. Following isolation, hippocampi were immediately snap-frozen by placing the sample tubes into dry ice and transferred to cold storage at -80 °C until sample processing.

### Nicotinamide treatment of HEK293T cells

HEK293T cells were cultured in Dulbecco’s Modified Eagle Medium + GlutaMAX™ (DMEM; catalog number 10566016, Gibco™, Thermo Fisher Scientific) supplemented with 1% Penicillin-streptomycin (10,000 IU/mL, catalog number 15140122, Thermo Fisher Scientific) and 10% fetal bovine serum (referred to as complete media). Cells were kept in a humidified incubator at 37 °C and 5% CO_2_ with media changed every 2-3 days until cells reached 80-90% confluency. Cells were collected by enzymatic dissociation, divided into 3 x 5 mL aliquots, and transferred to 15 cm^2^ cell culture dishes, with an additional 10 mL of media added to each dish following cell plating. Cells were cultured until they reached 50-60% confluency before undergoing treatment with the pan-sirtuin inhibitor, nicotinamide (NAM). Media was aspirated from each culture dish and replaced with fresh complete media (0 mM NAM), or complete media supplemented with 3 mM or 10 mM NAM (Millipore Sigma, catalog number N0636). Cells were cultured for a further 18 h and collected by enzymatic dissociation. Cells were washed twice with DPBS before proceeding to nuclei isolation and histone extraction as described in SI Methods (Histone Extraction).

### MS sample preparation

We evaluated two different workflows against the established Trypsin with peptide N-terminal and K derivatization protocol: denaturing vs. non-denaturing and derivatized vs. non-derivatized for Arg-C Ultra (MS grade, Cat number: VA1831, Promega™) and r-Chymotrypsin (rChymoselect, MS grade, Cat Number: CS3332042, Promega™; Table S1 and Figure 1). We also introduced TMT-labeling of Arg-C Ultra and r-Chymotrypsin digested peptides as an alternative to derivatization with propionic anhydride.

#### Protease digestion

Extracted histone samples were subjected to proteolytic digest with Arg-C Ultra or r-Chymotrypsin in 100 mM ammonium bicarbonate (AMBIC) under the conditions shown in Tables S1 and S2, according to the manufacturer’s recommendations. The pH of all buffers was tested using pH strips (Cat Number: 13640521, Fisher Scientific) to ensure the pH was maintained at 8-8.5. Digestions with Arg-C Ultra (1:100) and r-Chymotrypsin (1:40 or 1:10) were performed in 10 µL reactions at 37 °C and RT, respectively, for 2 h in a thermal cycler (Biorad, MJ Mini). For histones extracted from rat hippocampi, we used Arg-C Ultra (1:10) and r-Chymotrypsin (1:10) in 20 µL reactions. For histones extracted from HEK293T cells that were untreated or treated with NAM, we used 2.5 µg of histones per reaction (*n* = 3 digestion replicates/condition) digested with Arg-C Ultra (1:50) and r-Chymotrypsin (1:10) in 20 µL with 100 mM TEAB pH 8.5 as the digestion buffer. When labeling with TMT, HEK293T histones (5 µg) were digested in 100 mM TEAB pH 8.5, with Arg-C Ultra (1:100) or r-Chymotrypsin (1:10).

#### Derivatization (Propionylation)

Samples belonging to the Trypsin digestion groups were propionylated prior to digestion to prevent cleavage at the C-terminal of unmodified K residues. Samples were suspended in 50 mM AMBIC pH 8.0 to a final volume of 20 µL. The propionylation reagent was prepared as previously described ^1^ and all propionylation steps were performed in a fume hood. One batch of proprionylation reagent was used for up to 4 samples, and the propionylation was performed twice. Following the second round of propionylation, samples were dried in a vacuum centrifuge and stored at -80 °C until proteolytic digestion. Dried samples were resuspended in 10 µL digestion buffer containing Trypsin (Trypsin Gold, MS Grade, Promega, V5280) in 100 mM AMBIC pH 8.5. For the ‘Trypsin + Urea’ group, the digestion buffer also contained 2 M Urea and 5 mM TCEP (Table S2). Digestion reactions were incubated at 37 °C in a Thermomixer (Eppendorf) for 6 h. The reaction was stopped by freezing at -80 °C. The next day samples were thawed and dried in a vacuum concentrator. Two more rounds of propionylation were performed on Trypsin-digested samples and samples digested with Arg-C Ultra and r-Chymotrypsin (volumes adjusted with ddH_2_O to final concentration 50 mM AMBIC) to label peptide N-termini and remaining K residues (Table 1). Finally, propionylated peptides were dried and resuspended in 10 µL 0.1% formic acid (FA) in de-ionized water.

#### TMT labeling

Histone peptides suspended in ddH_2_O were mixed with TEAB to a final concentration of 100 mM, pH 8.0. Arg-C Ultra-generated peptides were labeled with TMT^10^-126, and r-Chymotrypsin generated peptides were labeled with TMT^10^-131 (cat no 90309, Thermo Scientific; monoisotopic mass = 229.162932) as per the manufacturer’s instructions (peptide:TMT ratio 1:8, final concentration of anhydrous acetonitrile = 44%) for 1 h at RT in a 9 µL reaction. Free TMT was quenched by adding 1 µL of 5% hydroxylamine to the reaction and incubating for 15 min at RT. Labeled peptides were dried in a vacuum concentrator and resuspended in 0.1% FA in de-ionized water.

Peptide concentration was determined in samples digested with Trypsin (non-denaturing conditions) by colorimetric peptide assay (Pierce Colorimetric Peptide Assay Kit, Thermo Scientific). Peptide samples were diluted 1:100 with 0.1% FA, and 20 µL (∼ 50 ng) aliquots were loaded onto Evotips (Evosep) following the manufacturer’s instructions.

### LC-MS/MS

Mass spectrometry analysis was performed using a Thermofisher Scientific Fusion Lumos Tribrid Mass Spectrometer configured with an electrospray ionization (ESI) source and operated in positive ion mode. The instrument was interfaced with an Evosep One nanoLC system (Evosep). The mobile phase comprised Solvent A (H_2_O with 0.1% FA) and Solvent B (ACN with 0.1% FA) (LC-MS grade, Fisher Scientific).

Reversed-phase HPLC separation was achieved using a custom-packed analytical capillary column (25 cm length, 150 nm internal diameter) containing Waters BEH C18 resin (1.7 µm particle size). Eluted peptides were introduced into the mass spectrometer via nanoelectrospray with a 2 kV spray voltage applied to the column inlet. Peptide fragmentation was performed using High-Energy Collisional Dissociation (HCD) in the Orbitrap. For non-TMT peptides, a fixed collision energy of 30% was applied, while TMT-labeled peptides were fragmented using a stepped normalized collision energy (NCE) of 30%, 40%, and 50% ^23^. The analytical method employed a 15 spd LC gradient (88 minutes) at 220 nL/min.

#### DDA

Full MS scans were collected at 120K resolution in the Orbitrap over a scan range of 375 – 1500 *m*/*z* in profile mode. Default charge state was set to +2, cycle time was 3 s, and maximum injection time was 50 ms. Included charge states were +2 to +7, dynamic exclusion was set to 5 s, and precursor mass tolerance was set to 10 ppm. All precursors above the minimum intensity of 5e^4^ during the 3 s cycle time, or up to the AGC target of 4e^5^ ions, were selected for HCD MS/MS scans in the Orbitrap at 7.5K resolution and collected as centroided data. Maximum injection time for MS/MS scans was set to 100 ms, with an AGC target of 5e^4^. Isolation was performed in the quadrupole with an isolation window of 1.6 *m*/*z*.

### Data analysis

#### Protein and PTM identification

MS raw files were processed in FragPipe (v24.0) following the recommended guidelines for the HiP-Frag workflow, with some modifications ^20^. Data were searched against a restricted database containing extracted human or rat histone sequences, contaminants and decoys (*Homo sapiens*: 342 sequences, 171 decoys; *Rattus norvegicus*: 292 entries, 146 decoys; contaminants lists were derived from and curated by Cambridge Centre for Proteomics (CCP) cRAP) with 1% FDR at the peptide and PSM level. The enzyme cleavage parameters were adjusted for Arg-C Ultra and r-Chymotrypsin to cleave after R or FYLM (not before P), respectively. Up to 2 missed cleavages were allowed for Arg-C Ultra and Trypsin (set to cleave after R only) and up to 3 missed cleavages were allowed for r-Chymotrypsin. An additional Trypsin search was conducted using the default cleavage specificity (KR) and missed cleavages (5) as an added assessment of the effects on peptide diversity resulting from incomplete propionylation of internal K residues. N-terminal propionylation was enabled as a static modification for all propionylated samples. For propionylated and non-propionylated samples, propionylation was set as a variable modification on K to account for endogenous propionylation in the case of the latter ^24^. Lists of variable modifications and detailed mass offsets are provided in SI Table S1. For TMT-labeled samples, the monoisotopic mass of the intact label (+229.162932 Da) was set as a static modification on peptide N-termini and as a variable modification on K, and all other PTM declarations were consistent with unlabeled samples. Label-free quantification (LFQ) and match-between-runs (MBR) were enabled. The propionyl and TMT masses were stripped from labeled peptidoforms during data processing to enable direct qualitative comparison of identified peptidoforms between labeled and unlabeled conditions.

#### Data processing and statistical methods

Data resulting from HiP-Frag output were imported into RStudio (RStudio 2025.09.2+, Build 418) and analyzed using custom R scripts. Reproducibility was assessed by calculating coefficients of variation (CV) from log2-transformed peptide intensities across technical replicates (*n* = 4 per condition for HEK293T samples; *n* = 5 for rat hippocampal samples). Digestion efficiency was evaluated by quantifying the proportion of peptides with 0, 1, or ≥2 missed cleavages. Labeling efficiency was calculated for both propionylation and TMT derivatization using two complementary metrics. For each peptide, lysine residues were classified based on modification status: labeled (bearing the expected derivatization mass: +56.026 Da for propionyl; +229.163 Da for TMT), free (unmodified), or biologically modified (e.g., acetylation, methylation). Biologically modified sites were excluded from calculations as they are not substrates for chemical derivatization. Site-based efficiency was calculated as labeled sites divided by the sum of labeled and free sites, expressed as a percentage (Efficiency (by site) = [labeled sites / (labeled sites + free sites)] × 100). Intensity-weighted efficiency was calculated by weighting each site count by peptide MS1 intensity, accounting for relative peptide abundance (Efficiency (by intensity) = [Σ(labeled sites × intensity) / (Σ(labeled sites × intensity) + Σ(free sites × intensity))] × 100). For TMT samples and propionylation, only lysine residues were considered. Informative peptides were defined as those that were fully labeled, contained ≤1 missed cleavage, and were detected in ≥3 replicates. Histone protein sequence coverage was calculated as the percentage of theoretical amino acid sequence represented by identified peptides. PTM diversity was quantified by counting unique modification types at specific residue positions (e.g., H4 K8ac K12ac = 2 acetylations). For histones extracted from frozen-thawed rat hippocampal sections processed using the RIPUP protocol, peptidoforms were retained if detected in ≥2 replicates. Data are presented as means ± standard deviation or medians where appropriate. Data were imported into Skyline (v 26.1.0.057 (c07debd50)) for visualization of chromatographic peaks corresponding to peptidoforms of interest.

#### Histone PTM quantification from DDA-LFQ data

Peptidoform-level quantitation was performed in FragPipe by disabling ‘MaxLFQ’ and ‘normalize intensity across runs’ to preserve changes in PTM abundance due to NAM treatment. Searches were conducted independently for Arg-C Ultra and r-Chymotrypsin without chemical derivatization (unlabeled DDA-LFQ) allowing up to 3 missed cleavages. To maximize signal recovery, all artifact-bearing forms of each peptidoform (Met oxidation, dehydration) were collapsed to a single biological modification state by stripping artifact masses from the modified sequence string, then summing intensities across variants sharing the same peptidoform identity per sample. For histone variants producing near-identical peptide sequences at the same protein positions (e.g., H2B type 1-J and H2B type 3-B), intensities were summed after canonical mapping to avoid duplicate quantitation of the same modification site.

For each histone (H2A, H2B, H3, H4, and H1 variants), per-sample total intensities were scaled to the grand mean across all samples within each histone, correcting for loading differences while preserving treatment-induced changes in individual peptidoforms. Intensities were log_2_-transformed, and missing values were imputed using k-nearest neighbors (kNN, k = 10; Bioconductor ‘impute’ package), restricted to dose groups where at least 2 of 3 replicates had measured values. Groups with 0 or 1 measured replicates were left as missing, ensuring that kNN was applied only to genuine single-replicate gaps arising from stochastic DDA sampling (missing at random). Peptidoforms with complete data across all dose groups after imputation entered the quantitative analysis; those with remaining missing groups were reported separately as qualitative presence/absence observations. Differential abundance was assessed using the ‘limma’ package with empirical Bayes moderated t-statistics. A cell-means model was fitted with NAM dose (0, 3, 10 mM) as a factor, and contrasts were tested for 3 mM vs 0 mM and 10 mM vs. 0 mM. P-values were adjusted for multiple testing using the Benjamini–Hochberg procedure. Peptidoforms are reported at a significance threshold of adjusted *p* < 0.05.

Dose-response concordance of peptidoform-level quantitative changes was assessed by plotting limma-derived log_2_ fold-changes for the 3 mM vs 0 mM and 10 mM vs 0 mM NAM contrasts against each other. Peptidoforms carrying acetylation or acylation modifications at known sirtuin-target sites were annotated to evaluate whether observed changes were consistent with pan-sirtuin inhibition. Peptidoforms not detected in all three dose groups were excluded from moderated statistical testing and visualized separately as per-replicate intensity dot plots.

## Results

### Protease and digestion conditions comparison

To assess potential improvements over the conventional Trypsin digestion with propionylation derivatization protocol, we systematically evaluated two alternative proteases, Arg-C Ultra and r-Chymotrypsin, under various experimental conditions including propionylation (Prop), urea denaturation, and TMT as an alternative to propionylation labeling. Our analysis focused on key performance metrics including total peptide identifications, digestion efficiency, sequence coverage, and the generation of informative peptides suitable for PTM identification and quantification. The reproducibility of protease activity and subsequent analyses via the developed pipeline were found to be robust, with median peptide CVs < 5% for all conditions (Figure 2A).

**Figure 2.**
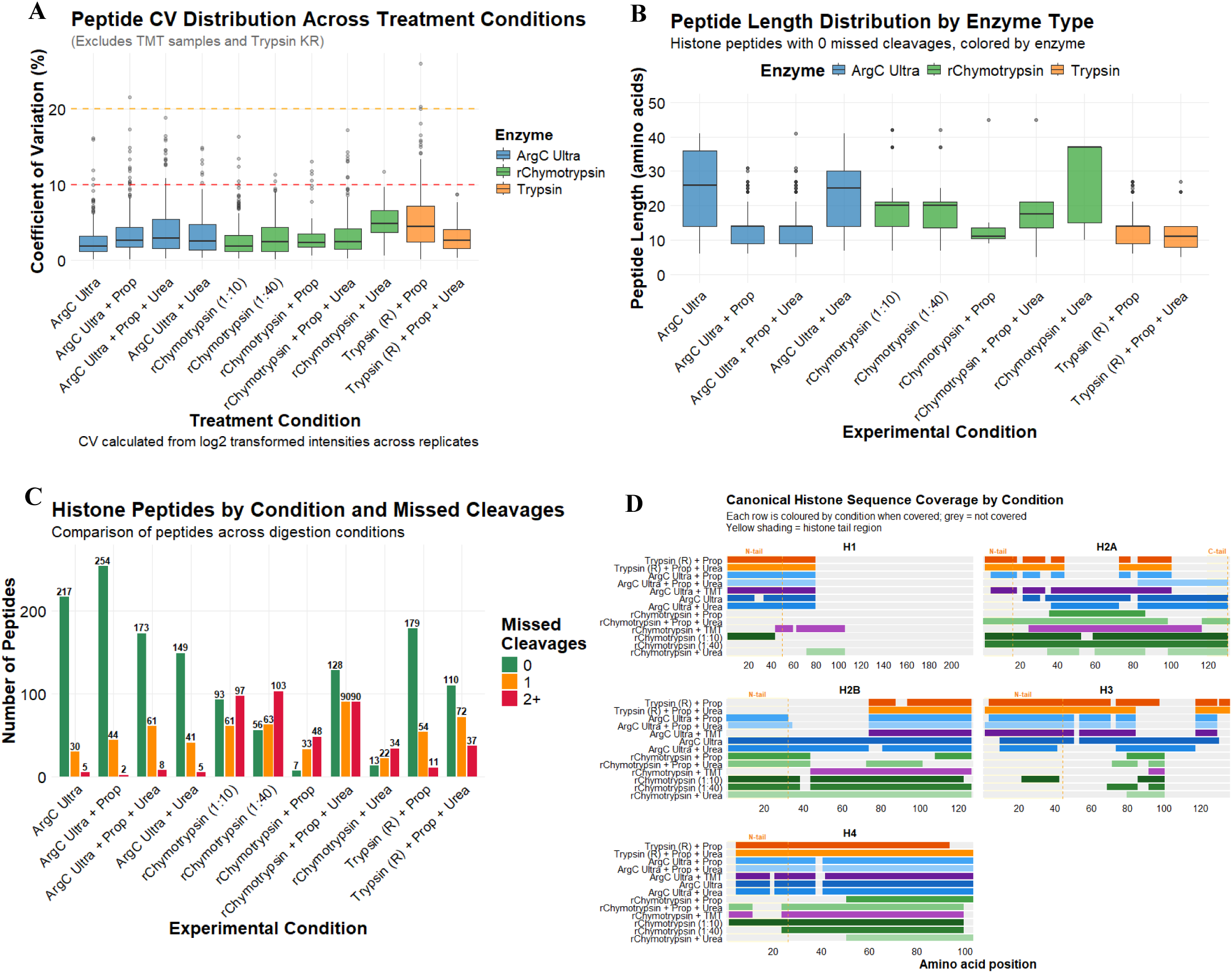
Comparative analysis of enzymatic digestion and propionylation strategies for histone post-translational modification detection in HEK293T cells (*n* = 4/condition). (A) Peptide coefficients of variation (CV) by enzyme type. All proteases yield CVs < 10 %. (B) Peptide length distributions for peptides with 0 missed cleavages. Propionylation facilitates identification of shorter peptides. (C) Number of histone peptides identified across experimental conditions, stratified by missed cleavages (0, 1, or 2+). Arg-C Ultra consistently yields the highest peptide counts with minimal missed cleavages. (D) Sequence coverage plots for canonical histone proteins across experimental conditions. H3 variants (H3.1, H3.2, H3.3) are aggregated due to high sequence similarity and canonical H1 represents H1.4. N-terminal and H2A C-terminal tail regions are indicated for canonical sequences.

The three proteases tested generated peptides with distinct length distributions when considering complete cleavage (Figure 2B). Short/hydrophilic peptides are often lost during chromatographic separation resulting from poor solid-phase retention, explaining over-representation of longer peptides (median peptide length >15 amino acids) when treating non-propionylated substrates with proteases. Further denaturation of substrates by urea increases the length range of identified peptides generated by r-Chymotrypsin, whose protease activity is known to be sensitive to any remnant secondary structures ^25^. In contrast, propionylation improves the hydrophobicity of short peptides, which reduces the median length of identified peptides in propionylated samples considerably, irrespective of urea denaturation. Trypsin and Arg-C Ultra showed similar cleavage specificity on propionylated histones, with a similar median peptide length of ∼15 amino acid in both conditions that was unaffected by the presence of urea.

Arg-C Ultra digestion yielded substantially higher peptide identifications than the conventional ‘Trypsin + Prop’ approach (Figure 2C). Digestion with Arg-C Ultra resulted in the identification of 217 distinct histone peptides with 0 missed cleavages. Additional propionylation improved peptide identification to 254 distinct histone peptides, likely due to increased hydrophobicity, peptide retention and separation during LC. Notably, Arg-C Ultra demonstrated superior digestion efficiency with most identified peptides showing no missed cleavage (∼84%, with and without propionylation, without urea), whereas ‘Trypsin + Prop’ produced a more heterogeneous mixture containing peptides with one or more missed cleavages. The addition of urea generally increased the number of missed cleavages across all enzyme types, though this effect was minimal with Arg-C Ultra (∼70% peptides without missed cleavage), and urea decreased the total number of peptides generated when the substrate was not propionylated. This could be attributed to the denaturing effect of urea being more pronounced on the protease activity than on the substrate that is already denatured by acid extraction.

r-Chymotrypsin followed similar trends, though fewer peptides were generated than Arg-C Ultra and higher missed cleavage rates were observed. r-Chymotrypsin performed similarly at the 1:10 and 1:40 enzyme-to-substrate ratios (90 and 56 peptides with no missed cleavages, respectively i.e., ∼20-25% peptides). In the presence of urea, fewer peptides were generated, though fully cleaved peptides still formed ∼21% of the total. However, in contrast to Arg-C Ultra, propionylation decreased the number of peptides generated with r-Chymotrypsin: only 7 fully cleaved peptides (∼8% peptides) were identified. In contrast to Arg-C Ultra, urea boosted overall digestion and fully cleaved peptide numbers of propionylated r-Chymotrypsin histone peptides to 128 (∼42% peptides), though substantially increasing missed cleavages.

### Histone protein sequence coverage

We assessed the impact of different enzymatic digestion strategies on histone protein sequence coverage, with histone sub-variants aggregated due to high sequence similarity (Figure 2D), while histone variants with significant sequence deviations were assessed for proteotypic coverage (SI Figure S1). Core histone H4 exhibited the highest sequence coverage across all conditions, with ∼75-95% coverage consistently achieved across most conditions. H3 coverage was highest (80%) with Arg-C Ultra and ‘Trypsin + Prop’, however sequence coverage provided by r-Chymotrypsin was low (∼10-20%). r-Chymotrypsin provided high sequence coverage for H2A (>95%), and exceptionally high proteotypic coverage for H2A.Z (72%, at both 1:10 and 1:40) compared to <6% coverage in propionylated conditions digested with Arg-C Ultra or Trypsin. r-Chymotrypsin provided the best proteotypic coverage of linker histone H1 variants (H1.2, H1.3, H1.5, H1.X), while Arg-C Ultra and Trypsin gave the highest coverage of H1.4 (canonical H1) (Figure 2D, SI Figure S1). The complementary coverage patterns observed across the three enzymatic approaches underscores the value of employing orthogonal digestion strategies to maximize histone proteome characterization and PTM site accessibility.

### Chemical rationale for TMT labeling

In this study, we used TMT labels as derivatization agents because of their high labeling efficiency and the chromatographic retention improvement they confer relative to underivatized peptides, independent of their conventional use as multiplexed isobaric labels for quantitating conditional changes in protein abundance ^17^. TMT and propionyl derivatization both target primary amines (N-termini and K ε-amines) via NHS ester and anhydride chemistry, respectively, forming stable amide bonds ^2^. However, the structural differences between these modifications impact peptide behavior during LC-MS/MS. Propionylation introduces a small aliphatic acyl group by nucleophilic attack of the peptide N-terminal and lysine ε-amine Nitrogen lone electron pair on a Carbonyl Carbon of the propionic anhydride structure (+56.026 Da), which increases peptide hydrophobicity due to the nonpolar ethyl chain of the propionyl moiety. This results in extended retention times on reversed-phase columns and can still result in co-elution of multiply modified histone peptides, particularly those derived from the K-rich N-terminal tails of histones H3 and H4. In contrast, TMT labels (∼229.1629 Da for TMT^10^) have multiple polar functional groups, including carbonyl oxygens and a tertiary amine within the reporter region. This structural composition confers moderate rather than high hydrophobicity, resulting in earlier retention times than propionylated counterparts (Figure 3A and 3B).

**Figure 3:**
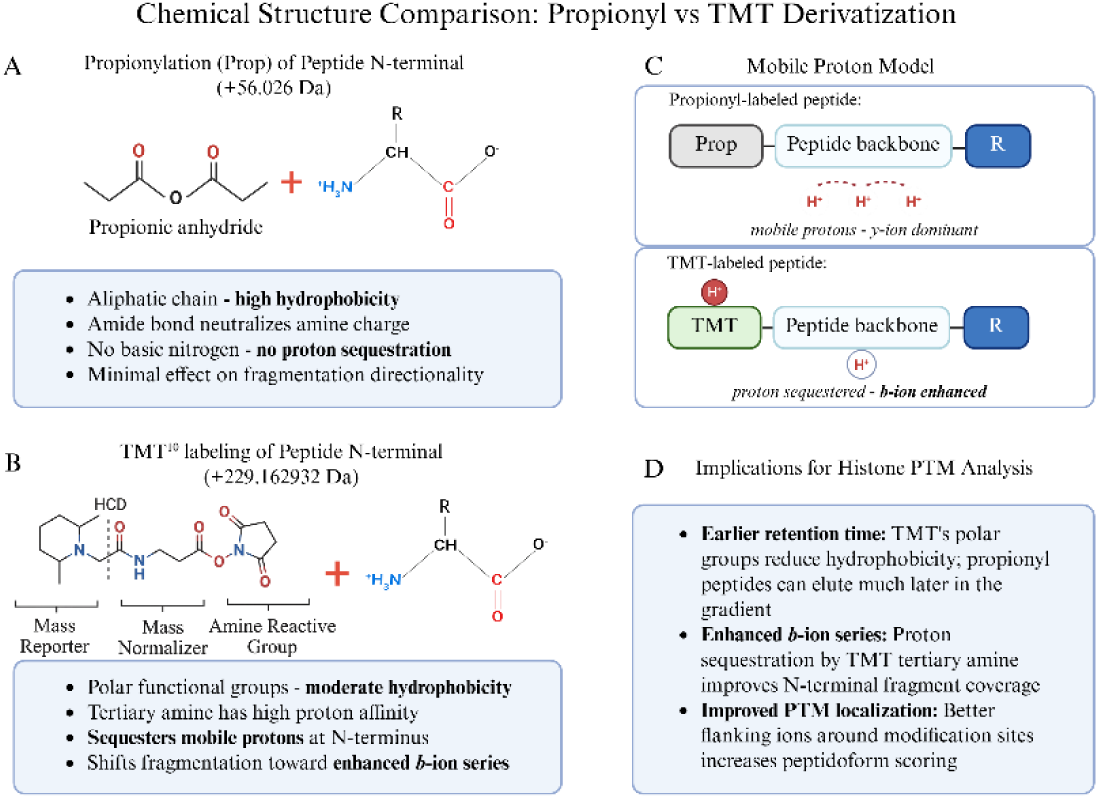
Chemical and structural comparison of propionylation and TMT derivatization strategies for histone peptide analysis. (A) Propionylation of peptide N-terminal amines using propionic anhydride (+56.026 Da). The propionyl group is a small aliphatic chain that increases hydrophobicity and neutralizes the amine charge but contains no basic nitrogen capable of proton sequestration. (B) TMT^10^ labeling of peptide N-terminal amines (+229.1629 Da). The TMT label comprises three functional regions: a mass reporter containing a tertiary amine with high proton affinity, a mass normalizer incorporating heavy isotopes (e.g., ¹³C, ¹⁵N), and an amine-reactive NHS ester group. The tertiary amine within the reporter region can sequester mobile protons during collision-induced dissociation. (C) Mobile proton model illustrating differential fragmentation behavior. In propionyl-labeled peptides, protons remain mobile along the peptide backbone, favoring *y*-ion formation. In TMT-labeled peptides, the tertiary amine sequesters a proton at the N-terminus, shifting fragmentation dynamics toward enhanced *b*-ion series generation. (D) Summary of implications for histone PTM analysis. Image of TMT structure in (B) was adapted from Thermo Scientific Pub. No. MAN0016969, Rev B.0, Pub. Part No. 2162457.5 (Figure 2). Created in https://BioRender.com.

Importantly, the tertiary amine within the TMT reporter region shows high proton affinity. According to the mobile proton model of peptide fragmentation ^26^, protons migrate along the peptide backbone during collision-induced dissociation and direct bond cleavage. In conventional tryptic and Arg-C-like peptides, mobile protons preferentially localize toward the C-terminus, favoring *y*-ion formation (Figure 3C) ^27^. The TMT tertiary amine sequesters a mobile proton at the N-terminal region of the peptide, shifting fragmentation dynamics to enhance *b*-ion series generation. This effect was particularly pronounced when using stepped collision energies of 30, 40, and 50 (normalized), which provided sufficient energy to fragment the peptide backbone while the TMT moiety retained the sequestered proton.

The enhanced *b*-ion coverage has direct implications for histone PTM analysis, where confident localization of PTMs requires the presence of flanking fragment ions on both sides of the modification site (Figure 3D). For histones, where multiple modifications often occur in close proximity on the same peptide, the enhancement of *b*-ions provided by TMT improves site localization confidence and peptidoform discrimination.

### Initial TMT assessment

Current methods of histone PTM analysis rely on efficient labeling/chemical propionylation of K and peptide N-termini to ensure robust PTM identification, quantification, and reliable comparison between two or more groups ^8,28,29^. However, it is well-documented that propionylation can be highly variable, with both under- and over-propionylation affecting downstream quantification ^9,12,28^. This was apparent when we included C-terminal of K as a potential site of Trypsin cleavage in the ‘Trypsin + Prop’ condition, where it is assumed that all lysines are propionylated and hence unavailable for Trypsin cleavage. Inclusion of K in cleavage specificity increased the number of identified peptides by ∼2.2 fold (∼2.4 fold considering peptides with 0 missed cleavages) compared to when only C-terminal of R Trypsin cleavage is included, suggesting that many lysines are not propionylated and therefore available for cleavage (SI Figure S2). For this reason and those described in the previous section, we performed an assessment of TMT as an alternative labeling strategy to propionylation. Although the use of TMT for histone PTM analysis has been suggested elsewhere ^30^ and demonstrated for histone H3 middle-down proteomics analysis ^31^, as well as for quantification of DNA damage-associated changes to histones following chromatin cross-linking in yeast ^32^, there has been no direct comparison of TMT to propionylation or other chemical derivatizing agents such as TMA ^10^, phenyl isocyanate (PIC) ^30^, or d^6^-acetic anhydride + PIC ^29^ for labeling histone-peptides. We compared the Arg-C Ultra and r-Chymotrypsin (1:10) generated peptides labeled with TMT to the established method of ‘Trypsin + Prop’, as these alternatives facilitate rapid sample processing with ≤ 2 h digestion times and only one round of labeling post-digestion. We first assessed the key metrics described in Figure 2 against ‘Trypsin + Prop’, labeling efficiency by site count and intensity, and then determined the number of informative peptides that could be used for quantification i.e., fully labeled peptides with ≤ 1 missed cleavage and no artifacts (e.g., Methionine oxidation). We did not perform a direct comparison to Trypsin labeled with TMT for two reasons: 1) TMT labeling of tryptic peptides requires both pre- and post-digestion labeling (as for propionylation), and unless sample cleanup (desalting) is performed prior to the second labeling, carryover of the label and/or hydroxylamine used to quench the labeling reaction would occur. This would introduce additional steps and variability into the workflow that would likely mitigate any benefit observed from the labeling approach itself thus rendering it an impractical choice, and 2) Arg-C Ultra cleavage should replicate the cleavage patterns of labeled Trypsin regardless, and as we have demonstrated in Figure 2, Arg-C Ultra performs exceptionally well with regard to cleavage efficiency and numbers of peptides identified compared to the conventional ‘Trypsin + Prop’ approach.

The median peptide CVs of TMT-labeled Arg-C Ultra and r-Chymotrypsin peptides were lower than ‘Trypsin + Prop’ peptides (Figure S2A). Median lengths of fully cleaved and TMT-labeled peptides generated by Arg-C Ultra and r-Chymotrypsin were similar to peptides from ‘Trypsin + Prop’ (∼15 amino acids; Figure S2B). TMT-labeling yielded ∼2.3-fold more fully cleaved peptides with Arg-C Ultra than the established ‘Trypsin + Prop’ approach (416 vs 179; Figure S1C); TMT-labeling of Arg-C Ultra yielded ∼1.6-fold more completely-cleaved peptides than ‘Arg-C Ultra + Prop’ (417 vs 254; Figure 2C and SI Figure S2C); TMT-labeling yielded >15-fold more completely-cleaved peptides than propionylation with r-Chymotrypsin (122 vs 7). Except for H4, histone sequence coverage decreased with Arg-C Ultra upon TMT-labeling in comparison to unlabeled (e.g., H3 decreased from 87% to 68%), although it was improved compared to Arg-C Ultra with propionylation (63%). Proteotypic histone sequence coverage remained similar or improved when r-Chymotrypsin peptides were TMT-labeled, except H2A.X (48% to 0%; Figure S2D). Artifact rates such as methionine oxidation and dehydration on S/E/T/Q were 2% and 4.5%, respectively, for ‘Trypsin + Prop’. This increased to 9.4% (dehydration) and 4.3% (oxidation) when Arg-C Ultra peptides were labeled with TMT, and both artifacts increased to 11.1% and 12.5%, respectively, in TMT-labeled peptides from r-Chymotrypsin digest (Figure S1E).

### Labeling efficiency of propionic anhydride vs TMT

Trypsin cleavage at non-propionylated K resulted in the loss of ∼58% of peptides (considering fully cleaved peptides) with ‘Trypsin + Prop’, which also prohibits the direct assessment of propionylation efficiency for Trypsin-digested samples (Figure S1F). However, we could directly assess propionylation efficiency for Arg-C Ultra and r-Chymotrypsin digests, where only one round of propionylation was performed post-digestion (Figure 4A). Considering only internal K as potential sites of propionylation (all peptide N-termini were considered propionylated, hence a static/fixed modification), propionylation efficiency calculated by site count (29-61%) was lower overall compared to estimation by intensity (33-71%). Efficiency of TMT-labeling was ∼92% by site count and ∼99% by intensity for peptides from Arg-C Ultra digest, and ∼91% by site count and ∼99% by intensity for peptides from r-Chymotrypsin digest.

**Figure 4.**
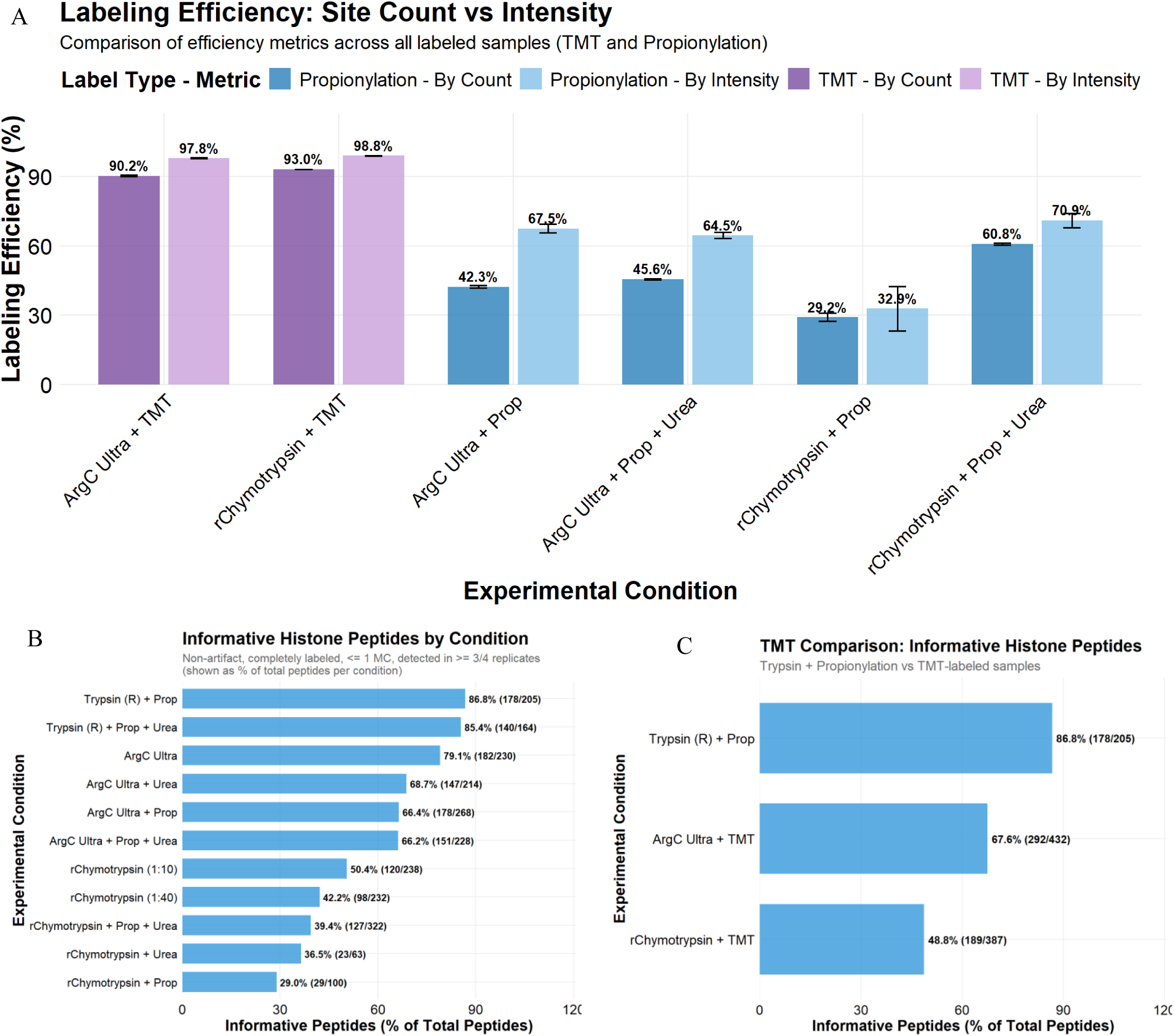
Comparison of internal lysine (K) labeling efficiency and informative peptide coverage across enzyme digestion strategies for histone PTM analysis. (A) Labeling efficiency metrics comparing propionylation and TMT-labeling approaches across all experimental conditions. Efficiency was calculated using two methods: by site count (darker bars) showing the percentage of sites that were successfully labeled out of all theoretically available sites for labeling, and by intensity (lighter bars) representing the sum of labeled peptide intensities divided by total peptide intensity. TMT-labeling efficiency by intensity was ∼98% in Arg-C Ultra and r-Chymotrypsin (1:10) digested samples, compared to propionylation efficiency by intensity of ∼68% in Arg-C Ultra and ∼33% in r-Chymotrypsin. Error bars represent standard deviation across *n* = 4 digestion replicates. (B) Percentage of IHP identified across different enzyme digestion conditions where IHP are defined as peptides containing completely labeled N-termini and lysine residues. Searching the Trypsin-digested samples with R cleavage specificity overestimates labeling efficiency, as any unlabeled K residues would result in cleavage at the lysine C-terminus. These peptides are not included in the search results, except in the case of a missed cleavage event. This is evident in Figure 4B, where ‘Trypsin + Prop’ and ‘Trypsin + Prop + Urea’ resulted in the highest IHP (86.8 and 85.4%, respectively), with unlabeled Arg-C Ultra the third highest (79.1%) but with the highest overall number of IHP (182). (C) TMT-labeling improved the number of IHP generated by Arg-C Ultra (292 vs 182) and r-Chymotrypsin (189 vs 120), which outperformed ‘Trypsin + Prop’ (178). Numbers in parentheses indicate IHP out of total peptides identified.

While the Garcia et al. ^1,2^ propionylation protocol was selected as the comparator given its dominant adoption in the field (291 citations), we note that the ammonium-containing buffers in this protocol introduce competing amine reactivity that reduces labeling efficiency relative to optimized approaches, and this issue was identified by Meert et al. (2015 and 2016) ^9,12^. The TMT advantage apparent in Figure 4A should therefore be interpreted as reflecting performance relative to the widely used protocol rather than against the best achievable propionylation efficiency.

Histone peptides that are reproducibly generated and identified with minimal variability are generally selected as the best candidates for precise quantification and highly confident comparative analysis, which we termed informative histone peptides (IHP). Building on established best practices for histone PTM data quality control ^29,33,34^, we defined these as peptides with ≤ 1 missed cleavage, identified in 3 out of 4 replicates, and lacking common artifacts like methionine oxidation or dehydration (Figure 4B). The effect of labeling and its efficiency was significant on IHP, with unlabeled samples outperforming propionylated samples overall when considering the proportion of IHP among total identified peptides. Although labeling (TMT and propionylation) and the presence of urea generally increased the total number of identified peptides, they slightly decreased the proportion of IHP. As mentioned above, since we substantially underestimate the total peptides in ‘Trypsin + Prop’ samples due to partial propionylation of K resulting in Trypsin-cleavage at these unlabeled K residues, we substantially overestimate the proportion of IHP (85-87%) in ‘Trypsin + Prop’ conditions. Arg-C Ultra digests provide a high proportion of IHP (∼80%) and r-Chymotrypsin digests yield ∼40-50% IHP, which decreases to ∼30-40% in the presence of urea or propionylation. TMT-labeling improved the number of IHP for both Arg-C Ultra and r-Chymotrypsin digestions (182 vs 292 and 120 vs 189, respectively), which outperformed ‘Trypsin + Prop’ and ‘Arg-C Ultra + Prop’ (both 178).

To further explore the impact of suboptimal labeling on the number of IHP, we compared our search results from Trypsin with cleavage specificity set to R (for fully labeled K residues) to a separate search with cleavage specificity set to KR (for partially/unlabeled K residues; SI Figure S2F). The number of fully labeled and properly cleaved peptides in ‘Trypsin + Prop’ (R) (178) constitutes ∼33% of the total number of peptides obtained from this method (sum of all peptides from ‘Trypsin + Prop’ (KR)).

### Histone PTM identification

Arg-C Ultra and r-Chymotrypsin digestion achieved high PTM coverage, detecting up to ∼120 unique modifications and over 15 distinct PTM types, including acetylation, methylation (mono-, di-, and tri-methyl states), ubiquitination (GG and RGG depending on digestion strategy), and acylations such as crotonylation, lactylation, succinylation, and malonylation (Figure 5). Arg-C Ultra resulted in high PTM numbers under most conditions, while ‘r-Chymotrypsin + Prop + Urea’ outperformed all other r-Chymotrypsin conditions substantially. Coverage of the most prevalent modifications (acetylation, methylation, and phosphorylation) was achieved across all conditions. Endogenous propionylation and butyrylation were identified in unpropionylated Arg-C Ultra and r-Chymotrypsin digested peptides, which cannot be distinguished from peptides labeled with propionyl +/- mono-methylation when propionic anhydride is used as a derivatizing agent. TMT-labeled peptides from Arg-C Ultra and r-Chymotrypsin digestion achieved comparable PTM numbers to conventional ‘Trypsin + Prop’ methods (∼120 PTMs) and enhanced identification of specific modification-types, including pronounced enrichment of negatively charged acylation marks such as succinylation and glutarylation (Figure 5B). The TMT-labeling approach detected a greater diversity of PTM classes than propionylation while offering superior labeling efficiency. Propionylation enabled identification of glyceroyl sites across all three enzymes, minimally detected in TMT-labeled or unlabeled samples, whereas Arg-C Ultra +/- urea increased identification of ubiquitination remnants. As amine-reactive labels modify the N-terminal amino acid of the GG remnant, charge neutralization occurs during propionylation; the overall charge state of a propionylated vs. unpropionylated GG-containing peptide would be ∼+1 vs. ∼+3, respectively. Arg-C Ultra samples without propionylation are therefore more likely to be selected for fragmentation (≥+2 charge) and exhibit increased ionization efficiency for PTMs that suffer from charge neutralization during propionylation.

**Figure 5.**
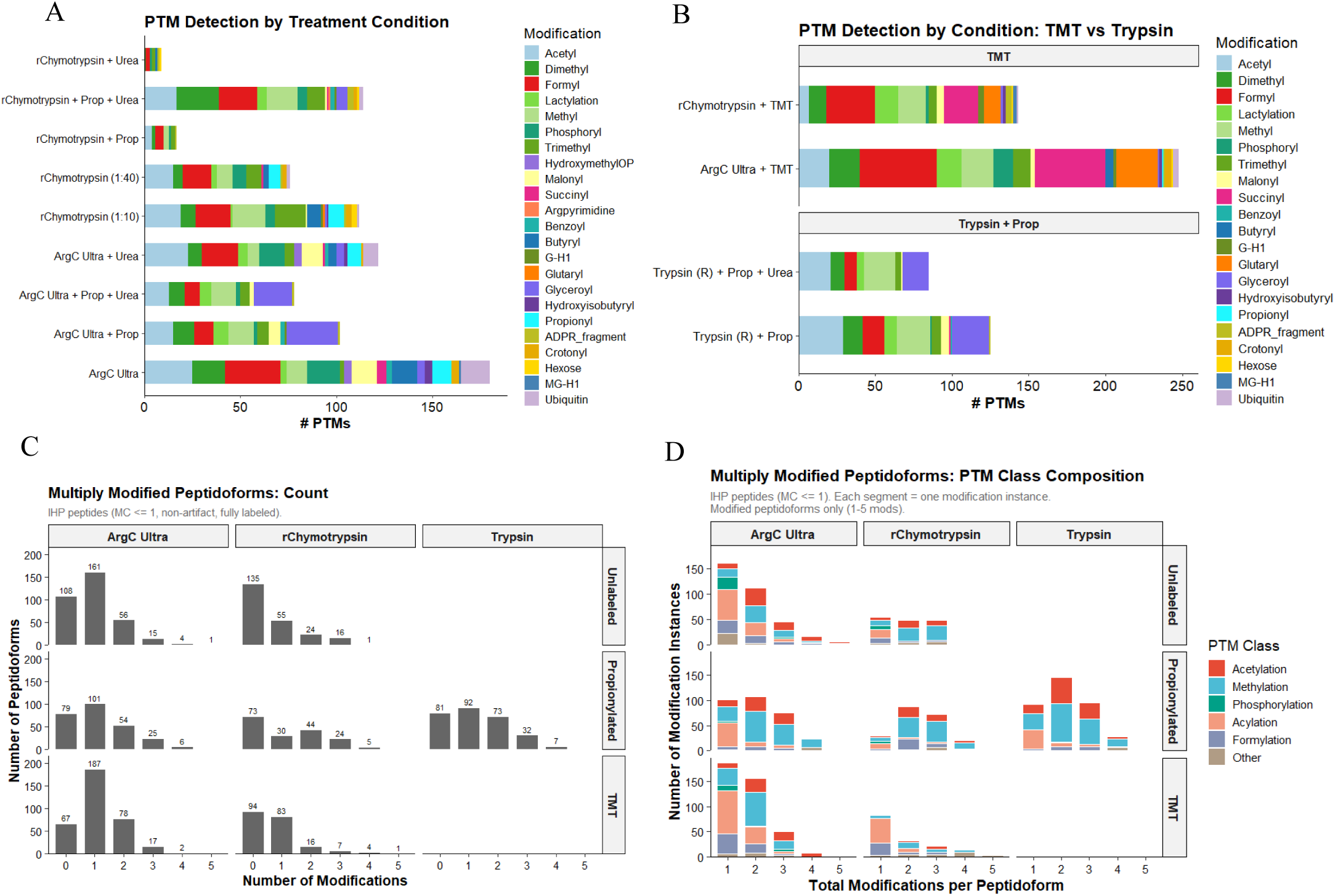
Diversity and abundance of histone post-translational modifications detected across enzyme digestion and chemical labeling strategies. (A) Total unique PTM sites identified per condition, stratified by modification type, across Arg-C Ultra, Trypsin, and r-Chymotrypsin digestion combined with propionylation (Prop), urea denaturation (Urea), Prop + Urea, or no derivatization (r-Chymotrypsin at 1:10 and 1:40 enzyme:substrate). (B) PTM identifications compared across TMT-labeled Arg-C Ultra and r-Chymotrypsin and propionylated tryptic peptides. Notable condition-specific detections: ubiquitin remnants and endogenous propionylation/butyrylation are identified in unlabeled Arg-C Ultra and r-Chymotrypsin digests; propionylation detects glyceroylation absent from labeled samples; TMT improves succinylation and glutarylation identification. (C–D) Multiply modified peptidoforms across all enzymes. (C) Unlabeled and TMT-labeled Arg-C Ultra yielded the most peptidoforms with ≥1 modification; Trypsin identified more peptidoforms with ≥3 modifications; unlabeled Arg-C Ultra and r-Chymotrypsin + TMT each identified a single 5-modification peptidoform. (D) PTM classes in multiply modified peptidoforms, grouped as acylations (crotonyl, butyryl, lactyl, malonyl, succinyl, glutaryl, hydroxyisobutyryl, glyceroyl, benzoyl, formyl, pyruvoyl, hydroxymethyl-OP, propionyl) and other modifications (ubiquitin remnants, citrullination, G-H1, MG-H1, argpyrimidine, hexose, ADPR/ADPR fragment, carboxymethyl-Q).

Multiply modified peptidoforms are analytically and biologically important given the role of PTM crosstalk in chromatin dynamics, which can influence protein-protein interactions and the accessibility of modification machinery ^35,36^. We therefore quantified peptidoforms carrying 0-5 modifications per condition (Figure 5C, 5D). Propionylated Arg-C Ultra and Trypsin peptides showed similar distributions of modified peptidoforms overall, though Trypsin produced more doubly modified peptides and the highest number of triply modified peptidoforms (32 vs 25 for Arg-C Ultra + Prop). TMT-labeled and unlabeled Arg-C Ultra conditions yielded the most singly modified peptides and the broadest acylation diversity. For r-Chymotrypsin, propionylation nearly doubled the yield of multiply modified peptidoforms compared to unlabeled digests (44 and 24 peptidoforms with 2 and 3 modifications, respectively, vs 24 and 16 unlabeled), with TMT labeling shifting the profile toward singly modified, acylation-enriched peptides. The PTM diversity captured by r-Chymotrypsin likely reflects its distinct coverage of H2A and H2B sequences, resolving region-specific modifications inaccessible to arginine-directed cleavage strategies.^35,36^

### Enhanced identification of histone succinylation and glutarylation with TMT

Lysine succinylation and glutarylation are acidic acyl modifications that reverse the charge of modified residues from +1 to −1, destabilizing nucleosome structure and promoting chromatin accessibility ^37,38^. Both modifications are enriched at promoters of active genes, regulated by acetyltransferases (KAT2A, p300/CBP) and sirtuins (SIRT5, SIRT7), and linked to metabolic state through their acyl-CoA donors ^39–41^. Dysregulation of these modifications has been implicated in cancer, protein-protein and DNA-protein interactions, and defective DNA repair ^19^, yet their comprehensive profiling has been limited by suppressed ionization of acidic peptides in MS-based workflows. A striking and serendipitous finding from our evaluation was the dramatically enhanced identification of succinyl-lysine and glutaryl-lysine sites in TMT-labeled peptides (Figure 5B, 6, and SI Figures S3-S6). While our current work and the recent study by Ryzhaya et al.^10^ demonstrate that chemical derivatization at the peptide level improves histone PTM analysis, our data reveal that TMT provides unique advantages for identifying acidic acylations other than improved chromatographic retention alone.

**Figure 6:**
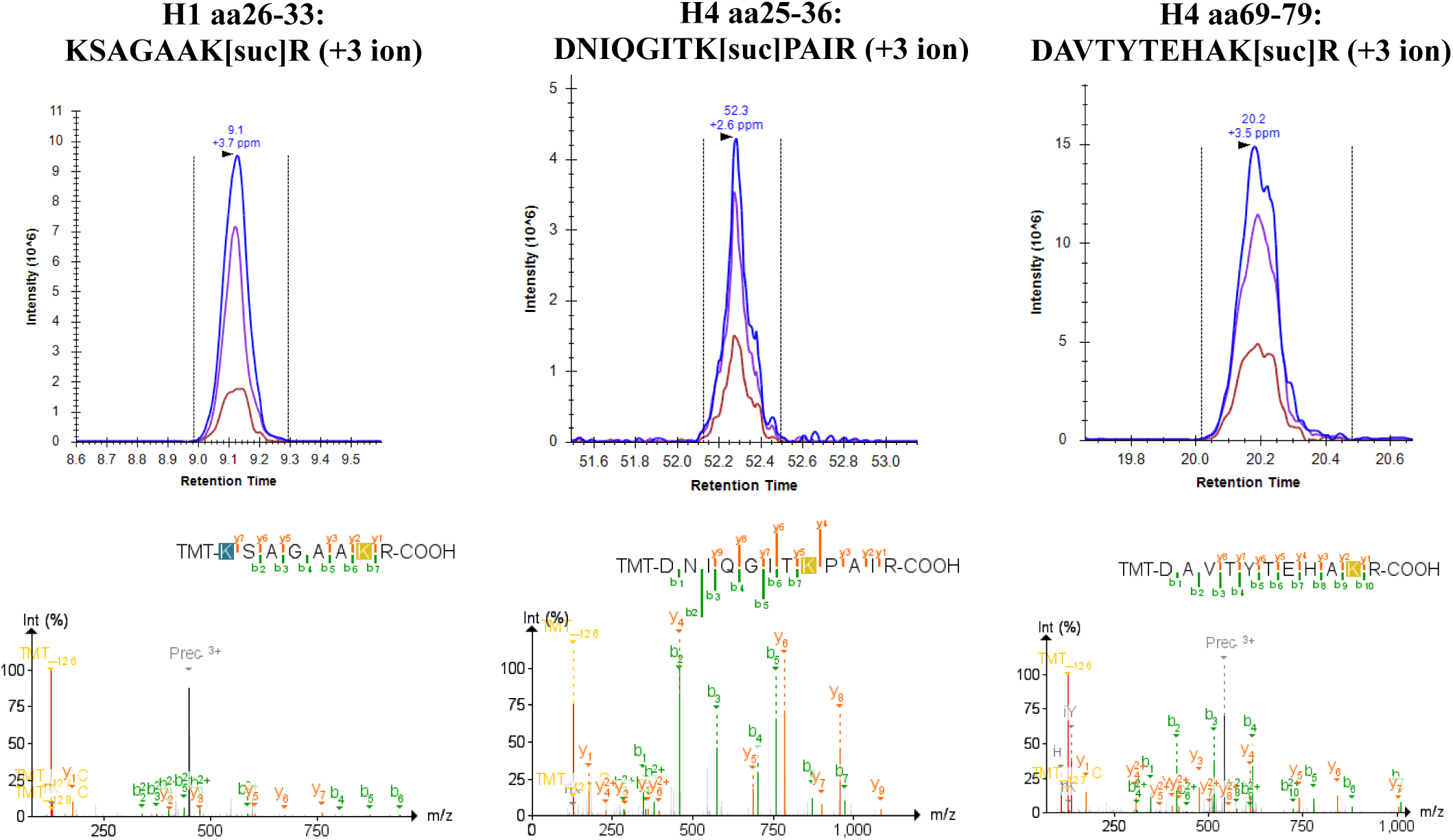
Extracted ion chromatograms (XICs) and corresponding annotated HCD MS/MS spectra of KSAGAAkR (+3 ion), DNIQGITkPAIR (+3 ion), and DAVTYTEHAkR (+3 ion) confirming succinyl-K localization in Arg-C Ultra digests labeled with TMT^10^. The modification site is unambiguously assigned based on continuous *b*-ion and/or *y*-ion series flanking the modified lysine residue. Precursor mass tolerance was within 10 ppm and fragment mass tolerance was within 20 ppm. Peak boundaries were confirmed to match the retention times reported in FragPipe.

Succinylation as a histone PTM is thought to occur less frequently than acetylation and methylation ^18,37^, though a recent study found that an increase in global histone succinylation is associated with longevity ^42^. In the current study, succinylation was often identified on first or second position K residues, adjacent K residues, or near the peptide C-terminus, which are challenging regions for confident PTM localization (Figure 6). For r-Chymotrypsin peptides labeled with TMT, succinylation sites were identified in H1.4 (9 sites), H2B (6 sites), H4 (3 sites), H2A variants (3 sites), and H3 (2 sites), including regions not covered by Arg-C Ultra or ‘Trypsin + Prop’ approaches (SI Figures S3 and S4).

The enhanced detection of succinyl-K with TMT is explained by the charge compensation provided by the tertiary amine within the TMT reporter region. Succinylation introduces a carboxylic acid moiety that impairs positive-mode ESI via charge state reduction. While propionylation increases peptide hydrophobicity, it neutralizes the charge of K residues and peptide N-termini. The TMT tertiary amine instead provides a compensating protonation site, rescuing ionization of succinylated peptides and increasing *b*-ion series flanking PTM sites (Figure 3 and Figure 6, SI Figures S3 and S4). This effect also extended to glutarylation (114.031694 Da) ^43,44^. Others have studied these negatively charged modifications using synthesized succinyl-K and glutaryl-K thioester derivatives in recombinant histone proteins ^19^. TMT-labeling thus offers a practical alternative for biological samples. The improvement in identification of succinylation (58 sites) and glutarylation (31 sites) with TMT-derivatization is attributable to improved ionization/fragmentation efficiency and PTM-site scoring, suggesting that succinylation of histone K residues may be more abundant than previously thought and revealing a PTM signature in HEK293T cells largely undetected by other methods and above what has been reported using MS-based methods ^18,37,42^.

### Quantitative analysis of histone peptides from RIPUP

NAM was used to induce global acetylation changes for evaluation of RIPUP quantitative performance. NAM is a potent pan-sirtuin inhibitor and is known to increase H3K9ac and H4K16ac ^45,46^. Conventional peptide-family ratio and site- occupancy approaches were unsuitable because NAM induced dose-dependent missed cleavage redistribution in Arg-C Ultra digests, evidenced by 259 peptidoforms detected exclusively in NAM-treated samples (Arg-C: 188; r-Chymotrypsin: 71), which would cause peptide family denominators to shift with treatment and artifactually suppress occupancy estimates. Instead, quantitation was performed directly on individual peptidoform intensities with histone-level loading correction, preserving 100% of robustly detected peptidoforms and enabling visualization of PTM redistribution across cleavage states. Unlabeled Arg-C Ultra identified 112 statistically significant peptidoform changes, 88 increasing with NAM dose in the expected direction. Hyperacetylation at sirtuin target sites is shown in Figure 7A: H3 peptidoforms carrying K9ac/K23ac reached significance at both doses across two backbone lengths (adj *p* < 0.001), alongside K14ac/K18ac/K23ac and K9ac/K14ac; on H4, K44ac increased at both doses, K12ac/K16ac and K16ac/K20me2 reached significance at 10 mM, and the tetra-acetylated K5ac/K8ac/K12ac/K16ac peptidoform was exclusive to NAM-treated samples. NAM also increased propionylation and butyrylation at H4K20 and propionylation at H2BK5, which are misassigned in propionylation workflows but readily resolved in RIPUP. Reciprocal changes in multiply modified peptidoforms, including K9me2/K23ac and K14ac/K18me2/K23ac, were also quantified, demonstrating RIPUP’s capacity for unbiased PTM crosstalk analysis (Figure 7B). A single succinylation site was detected but did not reach significance, consistent with our findings that TMT labeling is required to optimally ionize negatively charged acylations. While some increase in succinylation with NAM treatment might be expected given sirtuin involvement in histone desuccinylation ^47^, a recent study has suggested HDACs may play a more prominent role than sirtuins at these sites ^48^.

**Figure 7.**
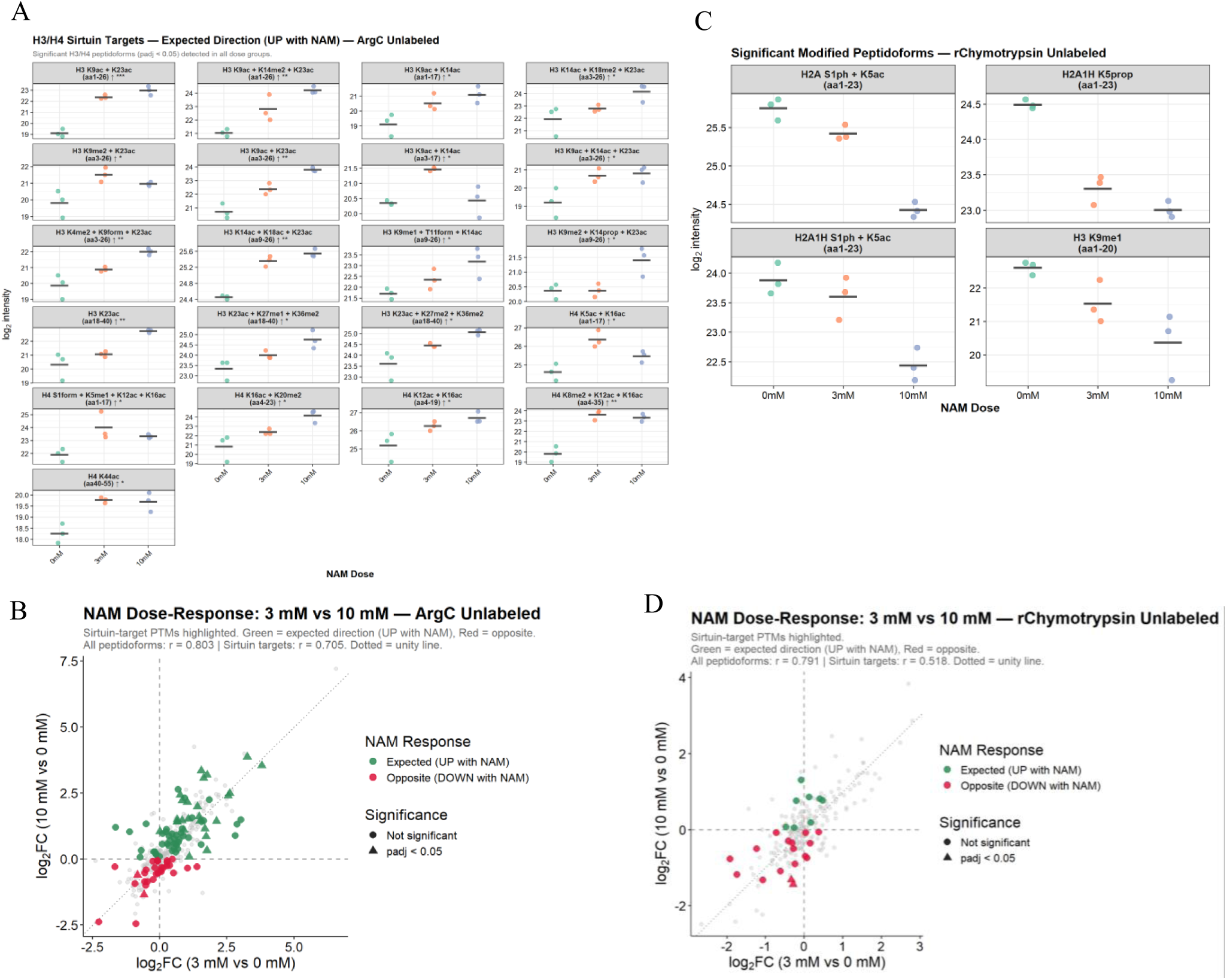
Quantitative histone PTM analysis in NAM-treated HEK293T cells with RIPUP. (A) Significant ArgC Ultra peptidoforms carrying sirtuin-targeted modifications (adj *p* < 0.05, detected across all dose groups). Arrows indicate direction; asterisks indicate significance (* adj *p* ≤ 0.05; ** adj *p* ≤ 0.01; BH-corrected). Peptidoforms were quantified by histone-level normalization, log_2_ transformation, kNN imputation (if detected in 2/3 replicates), and limma with pairwise contrasts. (B) Arg-C Ultra dose-response correlation of log2FC (3 mM vs 0 mM) against log2FC (10 mM vs 0 mM) for 342 peptidoforms (r = 0.803; sirtuin targets r = 0.705). Green: expected increase with NAM; red: opposite; triangles: adj *p* ≤ 0.05. (C) Significant r-Chymotrypsin modified peptidoforms (adj p < 0.05). Four modified peptidoforms reached significance, all decreasing with NAM dose. An additional 12 FDR-significant peptidoforms were unmodified, reflecting backbone redistribution from NAM-induced cleavage changes (not shown). (D) r-Chymotrypsin dose-response correlation for 235 peptidoforms (r = 0.791; sirtuin targets r = 0.518), confirming dose-dependent responses despite greater peptidoform diversity from FYLM cleavage specificity.

r-Chymotrypsin analysis quantified 235 peptidoforms across all dose groups, with 16 reaching FDR significance; 12 of these were unmodified peptidoforms reflecting backbone redistribution driven by NAM-induced cleavage efficiency changes. Among modified peptidoforms (Figure 7C), H2A S1ph/K5ac, H2A1H S1ph/K5ac, H2A1H K5prop, and H3 K9me1 all decreased significantly with NAM dose (adj *p* < 0.05), with the K9me1 decrease consistent with reciprocal K9 acetylation under sirtuin inhibition. An additional 52 modified peptidoforms were detected exclusively in NAM-treated samples, including succinylation and glutarylation on H1.4 and novel acetylation sites on H2A, H2A1H, and H3. Dose-response concordance was comparable to Arg-C Ultra (r = 0.791; Figure 7D). r-Chymotrypsin does not resolve H4 N-terminal tail peptidoforms carrying K16ac, underscoring the complementary coverage rationale of the dual protease RIPUP design.

We performed motif analysis to determine whether any specific flanking residues contributed to missed R cleavage (SI Figure S6 and S7), as this has not been previously reported for Arg-C Ultra or r-Chymotrypsin in the context of histone PTM analysis. Modifiable sites (K/R/STY) and basic residues (K/R/H) were enriched within 3 residues of missed R. Aspartic acid (D) was enriched at P1′ (Arg-C Ultra) and glutamic acid (E) was enriched at P2 (r-Chymotrypsin), suggesting that salt bridges may also contribute to cleavage site blocking, similar to their known effect on Trypsin cleavage efficiency ^49^.

### RIPUP of hippocampal sections

As a proof-of-principle experiment, we extracted histones from rat hippocampal sections and performed RIPUP using separate Arg-C Ultra and r-Chymotrypsin digestions. Total sample preparation time was ∼3 h and we identified 212 and 189 peptides with ≤1 missed cleavage with Arg-C Ultra (1:10) and r-Chymotrypsin (1:10), respectively (SI Figure S8A), with median CVs of ∼7% for both (SI Figure S8B). Combined, both enzymes detected 231 unique PTM sites, including all major PTM classes: acetylation, mono-, di-, and tri-methylation, phosphorylation, and ubiquitination (Figure 8A and B). Less common modifications were also detected, including endogenous propionylation and butyrylation. Biological monomethyls will be derivatized to butyryl and propionyl can also occur biologically, however these are indiscernible with propionylation-based protocols.

**Figure 8.**
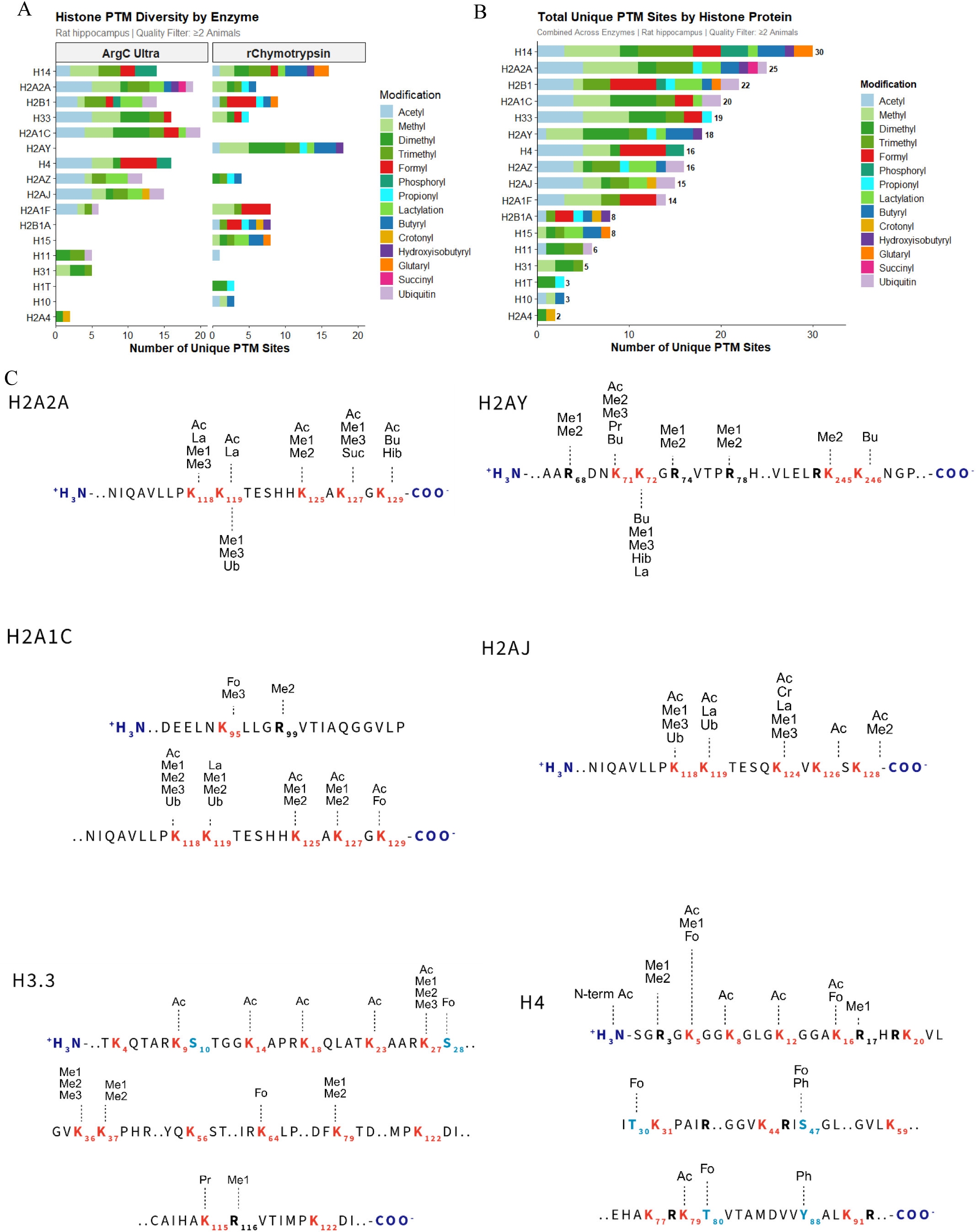
Comprehensive histone PTM landscape in rat hippocampus. (A) Histone PTM diversity detected per enzyme strategy in ≥2 animals. Stacked bars show unique PTM sites per histone using Arg-C Ultra (left) or r-Chymotrypsin (right), colored by modification type. (B) Total unique PTM sites per histone combining both enzymes; H1.4 and H2A2A showed the greatest diversity (30 and 25 unique sites, respectively). (C) Site-specific PTM maps for selected histone variants showing detected modification sites with K, R, S, and T residues highlighted. Subscript numbers indicate canonical residue positions. Abbreviations: Ac, acetylation; Me1/Me2/Me3, mono-/di-/trimethylation; Fo, formylation; Ph, phosphorylation; Ub, ubiquitination; La, lactylation; Bu, butyrylation; Cr, crotonylation; Hib, 2-hydroxyisobutyrylation; Pr, propionylation; Suc, succinylation; N-term, N-terminal modification. Data represent combined analysis from male rat hippocampi (n = 5). Figure 6C created in https://BioRender.com.

Analysis of selected core and variant histone sequences confirmed that highly modified N-terminal tails of H3 and H4 and ubiquitination sites K118/K119 on H2A were captured (Figure 8C). N-terminal acetylation and R3 mono- and di-methylation on H4, implicated in transcriptional regulation via PTM crosstalk ^50–52^, were identified, along with multiple peptidoforms of the H3 aa 9-17 and H4 aa 4-17 peptides enabling detection of combinatorial acetylation patterns with implications for chromatin structure and disease ^6,53–55^. The critical H3 region (aa 27-40) and K79 mono- and di-methylation were detected in Arg-C Ultra-digested samples; methylation of H3 K27, K36, and K37 functions as transcriptional activators or repressors via Polycomb silencing ^56–58^ and is crucial for interpreting gene expression anomalies in cancer ^59^.

RIPUP detected extensive PTM diversity on H1 and its variants. Canonical H1.4 contained 16 unique modification sites spanning residues 75-102, including succinylation at K75 and K90, crotonylation at K81, lactylation at K97, and arginine mono- and di-methylation at R79. H1.5 showed succinylation (K74) and crotonylation (K74, K80), while H1.9 displayed a complex modification profile at K96 (me3, pr, cr, bu) and K106 (me2, me3, cr, la, bu), plus ubiquitination (RGG remnant) at K97 by r-Chymotrypsin.

Core histone macro-H2A.1 exhibited arginine methylation at R69, R75, and R79, along with lysine modifications at K72 (ac, me2, me3, pr, bu) and K73 (me1, me3, cr, la, bu). H2B K59 emerged as a highly modified site, with seven distinct modification types at this single residue: acetylation, butyrylation, propionylation, lactylation, trimethylation, 2-hydroxyisobutyrylation, and ubiquitination. Glutarylation was also detected on H2A K119, demonstrating the benefit of retained positive charge on underivatized peptide N-termini for detecting negatively charged PTMs.

Formylation of K, S, T, and Y residues was a prominent feature in both HEK293T and rat hippocampal histones, an observation also made by authors of the HiP-Frag workflow across multiple cell lines ^20^. Although formylation can arise as a sample preparation artifact ^60^, this was not a feature of our workflow, and others have shown it to be an abundant and biologically important histone PTM on K ^61–63^.

## Discussion

### Computational stringency enables large-scale PTM discovery

*Positioning RIPUP within recent advances* Recent work by Vai et al. (2025) addressed the challenge of PTM confirmation at scale by developing HiP-Frag, integrating mass offset, open, and closed search strategies for confident novel PTM identification through rigorous computational filtering rather than comprehensive synthetic validation ^20^. Using this approach, they identified 60 previously unreported marks on core histones and 13 on linker histones across nine cancer cell lines and breast cancer tissue. Our study extends this framework by applying HiP-Frag principles to a systematic comparison of protease and labeling strategies. We demonstrate that the labeling strategy dramatically influences which PTM classes are detected: propionylation neutralizes positive charges on K residues without providing charge compensation for negatively charged modifications, whereas TMT’s tertiary amine rescues ionization of acidic acylations. This mechanistic explanation accounts for why succinylation and glutarylation may have been underreported with conventional workflows. Importantly, TMT and propionylation workflows reveal complementary rather than competing PTM landscapes. ^10^

Application of RIPUP to rat hippocampal tissue confirms that computationally driven PTM discovery extends to complex biological samples, consistent with Vai et al.’s analysis of breast cancer tissues ^20^. Detection of 231 unique PTM sites within a 3-hour workflow, including biologically critical marks such as H4 K5/K8/K12/K16 acetylation, H3 K27/K36/K37 methylation, and extensive linker histone modifications, validates RIPUP for time-sensitive biological discovery.

We applied 1% FDR at PSM and peptide levels, minimum detection in ≥3 digestion replicates (HEK293T) or ≥2 biological replicates (rat hippocampi), and CV filtering to ensure high-confidence assignments. The pragmatic validation approach used by Vai et al., which used targeted synthetic validation for ambiguous cases rather than comprehensive validation of all discoveries, is one we similarly adopt, and future studies could prioritize synthetic validation of the most abundant or biologically relevant succinylation and glutarylation sites identified here.

### RIPUP: A rapid multi-protease workflow for comprehensive histone PTM analysis

RIPUP’s dual-protease strategy maximizes histone sequence coverage within a minimal preparation window. Arg-C Ultra efficiently covers N-terminal tails and abundant PTM sites, while r-Chymotrypsin captures H2A variants, linker histone H1, and aromatic-rich regions inaccessible to arginine-specific cleavage, with overlapping coverage providing built-in orthogonal site validation ^64–66^. Application of the HiP-Frag framework identified 58 succinylation and 31 glutarylation sites in HEK293T cells, suggesting these acidic acylations have been systematically under-detected by conventional propionylation-based workflows. In rat hippocampal tissue, RIPUP identified 231 unique PTM sites across both enzymes within 3 h, including all major PTM classes, biologically critical marks on H3, H4, and H2A, and extensive H1 coverage largely inaccessible to Arg-C Ultra or Trypsin alone. This rapid turnaround is particularly advantageous in drug discovery and clinical contexts, such as intraoperative tumor assessment, where same-day epigenetic profiling has operational value.

RIPUP is not constrained by variability in chemical derivatization efficiency ^12,67,68^ and thus permits direct quantification of peptidoform abundance using established histone-level normalization methods ^29,33,34^. Our quantitative experiment (Figure 7) provides evidence that this approach is suitable and sensitive for detecting changes in peptidoform abundance in an *in vitro* model of hyperacetylation and facilitating PTM crosstalk analysis, while also revealing changes in propionylation and butyrylation that are normally misassigned in propionylation-based workflows. As the HiP-Frag workflow is designed for extensive PTM searches via detailed mass offsets list, the complete set of peptidoforms across multiple cleavage states can be included in quantitative analysis without filtering based on labeling efficiency. This ultimately minimizes experimental bias and improves quantitative accuracy. Although it was not performed here, DIA can be incorporated within the HiP-Frag framework using DDA spectral libraries generated by FragPipe/EasyPQP and DIA search with DIA-NN ^69^, which can further improve sensitivity and boost confidence in PTM identification. Moreover, it highlights important observations regarding protease kinetics and cleavage efficiency in a highly modified epigenetic landscape. To our knowledge, this is the first report of how histone PTMs at adjacent residues affect Arg-C Ultra and r-Chymotrypsin cleavage efficiency.

### Practical Considerations

The per-sample reagent cost of Arg-C Ultra is comparable to conventional Trypsin-based workflows. At a 1:100 enzyme-to-substrate ratio, Arg-C Ultra (Promega, $133/5 µg vial) yields approximately 100 reactions per vial at $1.33 per sample, compared to $0.69 per sample for Trypsin Gold MS-grade (Promega, $138/100 µg vial) at the 1:10 ratio recommended for histone digestion. This equates to a difference of $0.64 per sample. This marginal reagent cost is substantially offset by the elimination of two rounds of chemical derivatization required by propionylation-based workflows, reducing sample preparation to approximately 3 h. For multi-batch studies, clinical cohorts, or labile tissue samples where researcher time, sample throughput, and preservation of modification integrity are limiting factors, the RIPUP workflow offers a net reduction in total experimental cost. The r-Chymotrypsin protease used for orthogonal sequence coverage in the RIPUP workflow is available under the Early Access program (Promega). As a recombinant protease, production costs are expected to be comparable to existing MS-grade enzymes upon commercialization, though pricing has not been established. We have provided a cost analysis breakdown per sample for Trypsin, Arg-C Ultra, and the RIPUP dual protease workflow with and without labeling, which demonstrates the incorporating TMTzero as a derivatization agent only can significantly reduce per sample cost compared to TMT^10^ (SI Table S5). Reduction in hands-on researcher time by eliminating propionylation from the workflow brings the per sample cost down from ~$25/sample to ~$9/sample (unlabeled) and ~$16/sample for TMTzero-labeled samples.

### Limitations

We acknowledge that systematic comparisons were performed in a single cell line (HEK293T), and performance may vary with different sample types or extraction protocols, though broader applicability is demonstrated through hippocampal tissue analysis. Validation of the succinylation/glutarylation enhancement across additional cell types or primary tissues would strengthen these findings. Despite our dual protease strategy, sequence coverage gaps persist, particularly within globular domains. The reported PTM landscape reflects only regions accessible to the enzymes studied, though the RIPUP protocol is compatible with alternative digestion strategies to improve coverage ^64–66^.

PTM identification is controlled by the variable modification search space defined in the HiP-Frag workflow, and we found that the same limitations apply as per conventional closed searches in that the number of declared variable modifications should be limited to maximize results. Similarly, unanticipated modifications outside the variable modifications and detailed mass offset lists would not be detected, and the expanded search space increases false-positive risk despite FDR correction. Isobaric ambiguities such as trimethylation (+42.047 Da) vs acetylation (+42.011 Da) require high mass accuracy for confident discrimination, and some site assignments may remain probabilistic for peptides with multiple closely spaced modifiable residues. Notably, however, the RIPUP protocol is compatible with open-search pipelines for PTM discovery, so these constraints exist in downstream analysis rather than in the protocol itself.

Finally, endogenous propionylation and butyrylation cannot be distinguished from chemical derivatization artifacts in propionylated samples, a limitation the RIPUP workflow circumvents by omitting chemical derivatization, enabling quantitative comparisons of these modifications as we have demonstrated *in vitro* using NAM as a pan-sirtuin inhibitor.

## Conclusions

RIPUP is a rapid multi-protease workflow that reduces histone PTM sample preparation to 3 h while expanding PTM coverage through complementary protease strategies. Systematic evaluation across 40 samples and 10 conditions confirms Arg-C Ultra’s superior digestion specificity and demonstrates that TMT labeling offers a key advantage over propionylation-based derivatization: TMT’s tertiary amine provides charge compensation that rescues ionization of acidic acylations, revealing 58 succinylation and 31 glutarylation sites constituting an otherwise hidden ‘dark epigenome’. This suggests that the biological prevalence of negatively charged PTMs has been systematically underestimated by conventional workflows. Application to rat hippocampal tissue confirmed detection of 231 unique PTM sites within a three-hour preparation window.

For broad PTM discovery, particularly when acidic acylations are of interest, TMT-labeled Arg-C Ultra/r-Chymotrypsin is the recommended first-choice workflow, achieving PTM identifications exceeding the conventional ‘Trypsin + Prop’ benchmark while uniquely enabling ‘dark epigenome’ detection. For comprehensive sequence coverage across histone variants, the dual-protease RIPUP approach provides orthogonal coverage within a comparable timeframe. For cost-constrained applications focused on conventional PTMs, ‘Arg-C Ultra + Prop’ remains effective, with the caveat that negatively charged acylations will be under-represented. Collectively, RIPUP expands the analytical toolkit for high-throughput epigenetic profiling to accelerate mechanistic and therapeutic discovery.

## Supporting information

Supporting Information for Publication

## Supporting information

Supplementary methods (histone extraction, SDS-PAGE, FragPipe search settings); Figure S1, variant-specific proteotypic sequence coverage heatmap across conditions; Figure S2, quality metrics comparing TMT and propionylation workflows (CVs, peptide length distributions, missed cleavages, histone protein sequence coverage, chemical artifact rates, and cleavage specificity search parameters); Figure S3, H1.4 sequence coverage and succinylation sites in r-Chymotrypsin + TMT and Arg-C Ultra + TMT samples; Figure S4, representative extracted ion chromatogram and MS2 spectrum for succinylated H1.4 peptide KSAGAAKR in Arg-C Ultra + TMT samples; Figure S5, H4 sequence coverage and succinylation sites with representative fragment spectra in r-Chymotrypsin + TMT (A) and Arg-C Ultra + TMT (B) samples; Figure S6, Arg-C Ultra missed cleavage motif analysis from NAM-treated and untreated HEK293T histones; Figure S7, r-Chymotrypsin missed cleavage motif analysis from NAM-treated and untreated HEK293T histones; Figure S8, unique peptidoforms, missed cleavages, and CV distributions of rat hippocampal histone peptides obtained with the RIPUP protocol; Figure S9, SDS-PAGE gel images of histones extracted from HEK293T cell and rat hippocampi; Table S1, detailed FragPipe mass offset parameters; Table S2, protease digestion conditions; Table S3, denaturing conditions for histone digestion; Table S4, peptide lists of modified peptides identified in rat hippocampal sections using the RIPUP workflow (Arg-C Ultra and r-Chymotrypsin); Table S5, persample reagent and researcher time cost comparison.

## Data availability

The MS raw data files, annotations, Sample and Data Relationship Format (SDRF-Proteomics) ^70^, and FragPipe search results have been deposited to the ProteomeXchange Consortium (http://proteomecentral.proteomexchange.org) ^71,72^ via the PRIDE partner repository ^73^ with the dataset identifier PXD073683. The custom R scripts used for data analysis are available at: https://github.com/NataliePTurner/Histone-RIPUP.

## Acknowledgements

The authors would like to thank Dr Antonio Pinto in the Scripps Research Institute Multi-omics Core for consultation and technical insight. Support for this study was provided by National Institute on Alcohol Abuse and Alcoholism grants: T32 AA007456, AA013498 (MR, MB), P60 AA006420 (MR, JY), AA017447 (MR), AA029841 (MR), AA021491 (MR) and the Schimmel Family Chair.

## Notes

### Competing Interest Statement

The authors have declared no competing interest.

### Summary of Updates

Minor edits to methods to clarify data filtering and to the discussion to extend cost analysis discussion. Additional table added to supporting information on cost analysis (SI Table S5).

