## Supporting Information for Publication for "Rapid Histone Post-Translational Modification Analysis Using Alternative Proteases and Tandem Mass Tags"

#### Table of Contents

SI Methods....................S2

Histone extraction....................S2

SDS-PAGE....................S2

FragPipe Settings....................S3

References....................S3

**Figure S1.** Heatmap showing variant-specific proteotypic sequence coverage in each condition. ....................S4

**Figure S2.** Comprehensive quality metrics comparing TMT derivatization and propionylation workflows for histone peptide analysis....................S5

**Figure S3.** H1.4 sequence coverage and succinylation sites in r-Chymotrypsin + TMT and Arg-C Ultra + TMT samples....................S7

**Figure S4.** Representative extracted ion chromatogram (XIC) and matching mass spectrum for the H1.4 peptide KSAGAAKR (+3 ion) in Arg-C + TMT labeled samples. ....................S7

**Figure S5A.** H4 sequence coverage and succinylation sites in r-Chymotrypsin + TMT samples. ....................S8

**Figure S5B.** H4 sequence coverage and succinylation sites in Arg-C Ultra + TMT samples....................S8

**Figure S6.** Arg-C Ultra missed cleavage motif analysis from NAM-treated and untreated HEK293T histones. ....................S9

**Figure S7.** r-Chymotrypsin missed cleavage motif analysis from NAM-treated and untreated HEK293T histones....................S10

**Figure S8.** Peptidoforms, missed cleavages, and CV distributions of rat hippocampal histone peptides obtained with RIPUP. .....S11

**Figure S9.** SDS-PAGE protein profiles of histones extracted from HEK293T cells and rat hippocampal sections....................S11

**Table S1:** FragPipe Detailed Mass Offsets....................S13

**Table S2:** Protease digestion conditions for histones extracted from HEK293T cells....................S14

**Table S3:** Denaturing conditions for histones extracted from HEK293T cells....................S14

**Table S4:** Peptide lists of modified peptides identified in rat hippocampal sections using the RIPUP workflow (Arg-C Ultra and r-Chymotrypsin) ....................S15

**Table S5:** Per-Sample Reagent and Researcher Time Cost Comparison (5 µg Histone Peptide Input)....................S27

### SI Methods

#### Histone extraction

Histones were extracted from HEK293T cells using an established protocol as described by Sidoli *et al* (Sidoli et al., 2016) Briefly, frozen-thawed aliquots of HEK293T cells were pooled by transferring to a 50 mL falcon tube, and the tube topped with DPBS warmed to 37 °C. Cells were washed twice by centrifugation and the cell pellet was resuspended in chilled nuclear isolation buffer (NIB) with DTT and protease inhibitors (Pierce EDTA-free protease inhibitor cocktail, ThermoScientific). Cells were washed once in NIB, lysed with NIB containing 0.2% NP-40 alternative, and homogenized by gentle pipetting. The mixture was incubated on ice for 10 min and centrifuged to pellet nuclei. The nuclei pellet was washed three times with NIB by centrifugation to remove traces of detergent, and the supernatant was discarded after the final wash. For rat hippocampal sections, frozen samples were thawed on ice, washed once with chilled NIB, transferred to a 2 mL dounce homogenizer, and homogenized/lysed with 10 – 15 strokes of the fine pestle. All other nuclei isolation steps were the same as described for nuclei extraction from HEK293T cells.

Acid extraction of histones from the nuclei pellets was performed by resuspending nuclei in 0.2 M $H_2SO_4$ and incubating at 4 °C for 2-3 h with gentle rotation. Insoluble material was removed by dual rounds of centrifugation, and trichloroacetic acid (TCA; 100%) was added to the supernatant to a final concentration of 33%, vortexed to mix, and incubated on ice overnight to precipitate histones. Pelleting and washing of precipitated histones with acetone/HCl and 100% acetone were performed as previously described (Sidoli et al., 2016). The final histone pellets were resuspended in dd$H_2O$ and centrifuged briefly to pellet insoluble material. The supernatant was transferred to a new tube and assessed for quality by SDS-PAGE and protein concentration by BCA assay (Pierce™ BCA Protein Assay, cat number 23227, Thermo Scientific™). For HEK293T samples, the sample was divided into aliquots of ~5 µg protein according to Tables S1 and S2, with four technical replicates in each condition, resulting in a total of 40 samples. For rat hippocampal sections, 2.5 µg of purified histones were used per protease (5 µg total per sample).

#### SDS-PAGE

Aliquots of 5 µL extracted histones were mixed with 4x LDS sample buffer (NuPAGE™, Catalog number NP0007, Invitrogen™) and 10x sample reducing agent (NuPAGE™, Catalog number NP0009, Invitrogen™), to achieve a 1x final concentration in the sample. Samples were reduced for 10 min at 70 °C and 450 rpm in a Thermomixer (ThermoScientific). A volume equivalent to 2 µg total protein of purified histone H2A standard (Sigma-Aldrich, Cat number H9250, Sigma-Aldrich) was prepared in the same way. A 5 µL aliquot of protein ladder (BLUEstain™ 2 Protein ladder, 5-245 kDa, Cat Number: P008-500, Goldbio), samples, and H2A standard were separated by gel electrophoresis on a 4-12% Bis-Tris mini protein gel, 1.0–1.5 mm (NuPAGE™, Cat number: NP0321BOX, Invitrogen™) for 50 min at 150 V. The protein gel was placed into a clean gel tray and incubated with Acquastain gel stain (Bulldog Bio) to visualize protein bands. Gels were imaged on a gel imager with 5 s exposure (Azure biosystems c600).

#### FragPipe Settings

All default variable modifications and mass offsets were enabled as per default pipeline settings, except methylation (+14.01565 Da) was added as a variable modification to non-propionylated samples (default setting +70.0419 Da, the combined mass of a propionyl group and methyl group), and the +70.0419 Da mass was retained as endogenous butyrylation. Propionylation on serine, threonine, and tyrosine was removed from the detailed mass offset list for non-propionylated samples. For TMT-labeled samples, the monoisotopic mass of the intact label (+229.162932 Da) was set as a static modification on peptide N-termini and as a variable modification on K, and all other PTM declarations were consistent with unlabeled samples. Finally, there were specific mass shifts considered for Gly-Gly (GG; +114.0429 Da) or Arg-Gly-Gly (RGG; +270.1441 Da) remnants corresponding to cleavage of ubiquitinylated K (Arg-C Ultra and r-Chymotrypsin cleaved ubiquitin, respectively. These fragments can also be labeled with propionyl or TMT at the free amine at the N-terminus, so +170.0691 Da (GG + Prop) and +343.2058 Da (GG + TMT) for Trypsin or Arg-C Ultra-digested samples, and +326.1703 Da (RGG + Prop) and +499.307 Da (RGG + TMT) for r-Chymotrypsin-digested samples were included. Label-free quantification (LFQ) and match-between-runs (MBR) were enabled for quantification. The propionyl and TMT masses were removed from labeled peptidoforms during data analysis processing to enable direct qualitative comparison of identified peptidoforms between labeled and unlabeled conditions.

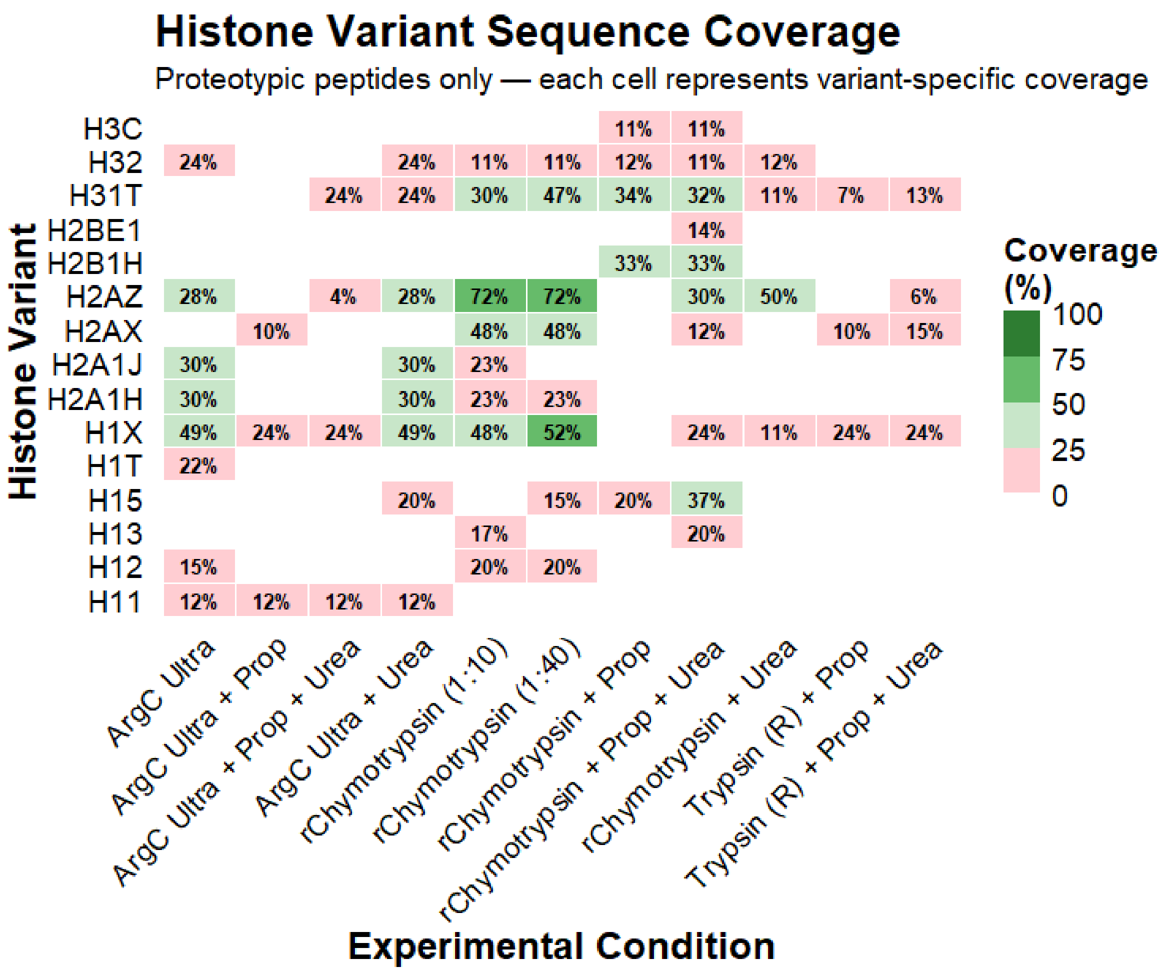


**Figure S1.** Heatmap showing variant-specific proteotypic sequence coverage in each condition.

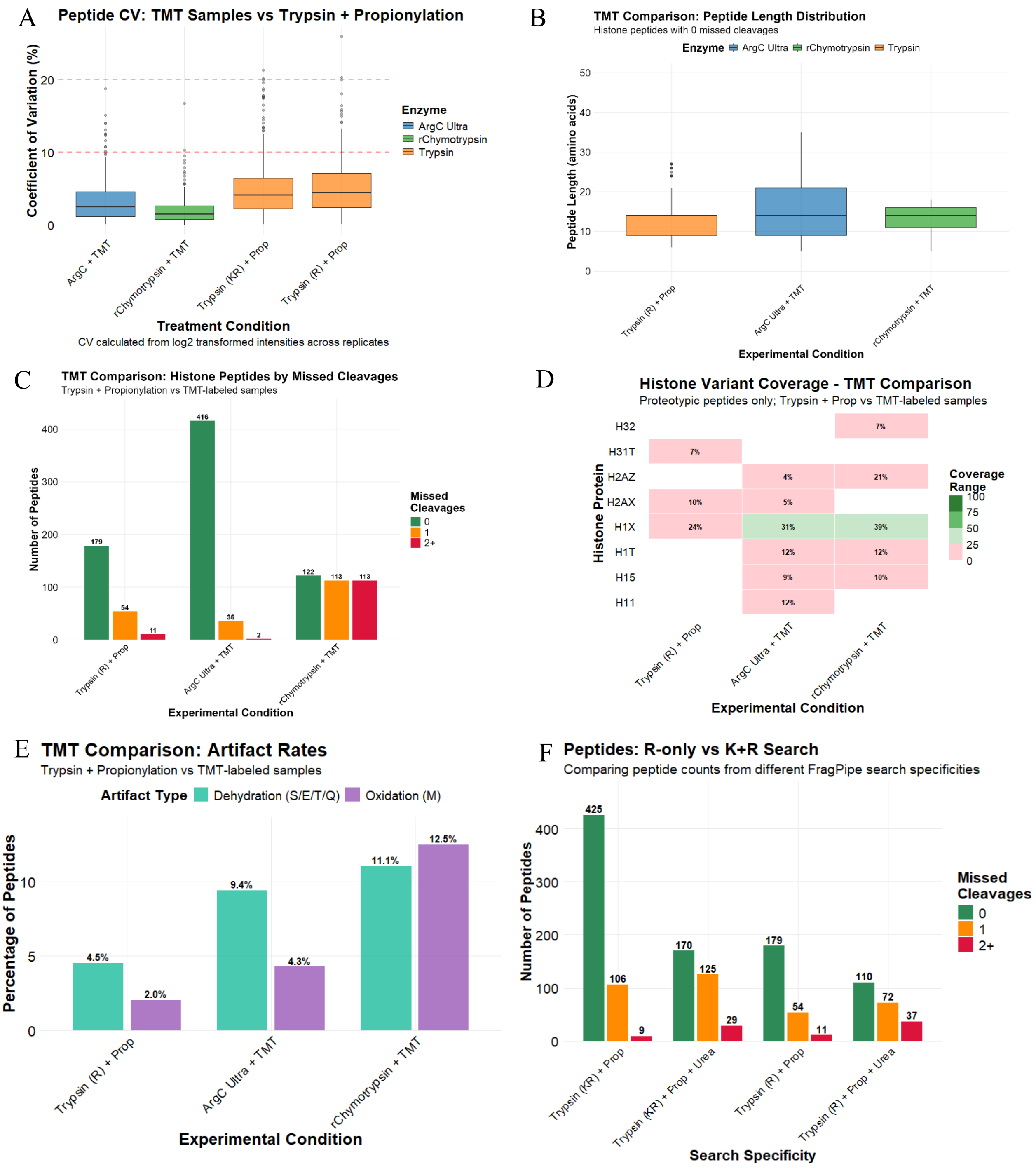


**Figure S2.** Comprehensive quality metrics comparing TMT derivatization and propionylation workflows for histone peptide analysis. **(A)** Coefficient of variation (CV) of peptide intensities across replicates. CVs were calculated from log2-transformed intensities. TMT-labeled samples (Arg-C Ultra, r-Chymotrypsin, Trypsin) demonstrate comparable or lower variability than Trypsin + Propionylation. Dashed lines indicate 10% and 20% CV thresholds. **(B)** Peptide length distribution for fully cleaved histone peptides (0 missed cleavages). Peptide length distributions are similar across Trypsin + prop and TMT-labeled samples (median ~15 amino acids). **(C)** Distribution of missed cleavages in identified histone peptides. **(D)** Histone protein sequence coverage heatmap using proteotypic peptides for histone variants. **(E)** Chemical artifact rates comparing derivatization methods.

Dehydration artifacts (S/E/T/Q) and methionine oxidation are shown as percentage of total peptides. **(F)** Effect of search enzyme specificity on peptide identifications. Comparison of R-only (fully labeled) versus K+R (partially labeled/unlabeled) search parameters across different sample preparation methods, stratified by missed cleavages.

H1.4 Succinylation sites: sequence coverage

Protease-specific site coverage

*r-Chymotrypsin + TMT*

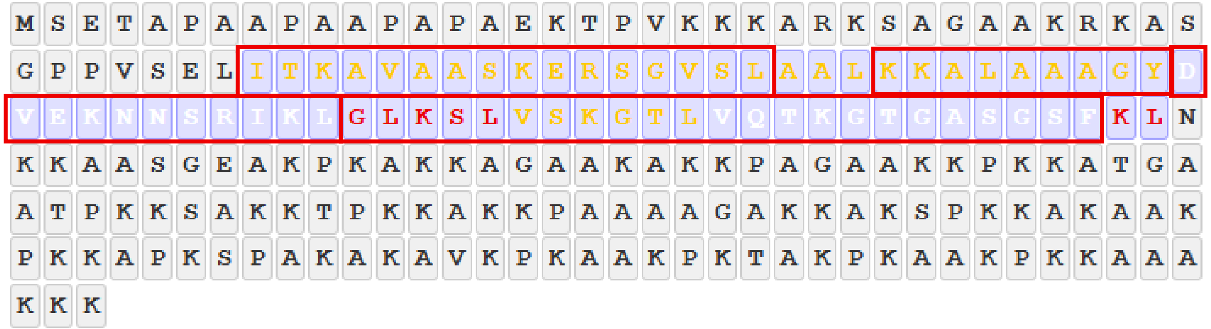


ITkAVAASKERSGVSL
ITKAVAASkERSGVSL
kKALAAAGY
KkALAAAGY
DVEkNNSRIKL
DVEKNNSRIkL
GLkSLVSKGTL
GLKSLVSkGTL; VSkGTL; VSkGTLVQTKGTGASGSF
VQTkGTGASGSF; VSKGTLVQTkGTGASGSF

*Arg-C Ultra + TMT*

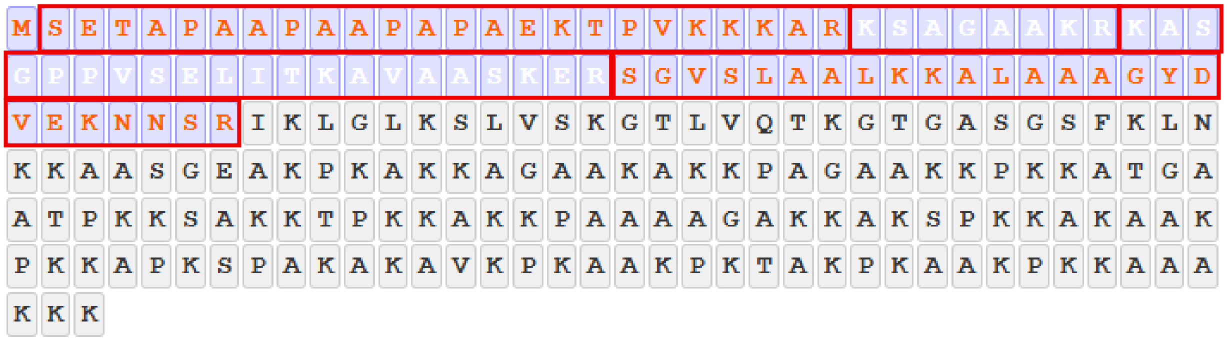


kSAGAAKR
KSAGAAkR
kASGPPVSELITKAVAASKER

KASGPPVSELITkAVAASKER
KASGPPVSELITKAVAASkER
SGVSLAALKkALAAAGYDVEKNNSR

**Figure S3.** H1.4 sequence coverage and succinylation sites in r-Chymotrypsin + TMT and Arg-C Ultra + TMT samples. Theoretical sequence coverage is indicated by purple shading. Red boxes indicate observed peptides/sequence coverage of succinyl-K peptides. Peptides highlighted in yellow indicate protease-specific succinyl-K site coverage.

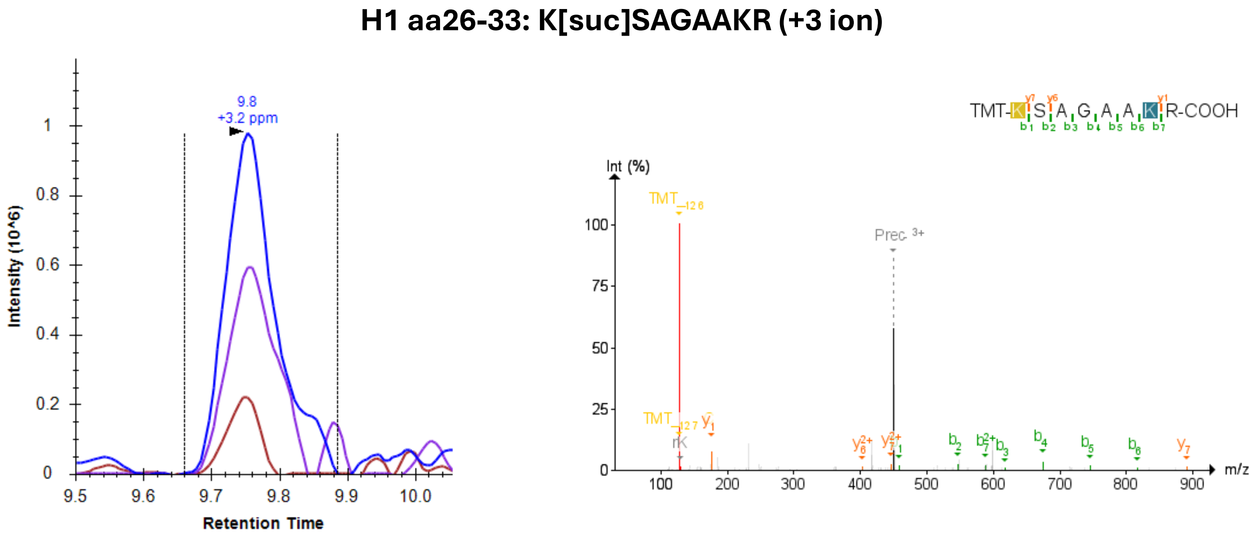


**Figure S4.** Representative extracted ion chromatogram (XIC) and matching mass spectrum for the H1.4 peptide KSAGAAKR (+3 ion) in Arg-C + TMT labeled samples. Peptide sequence with colors above the MS2 spectrum indicate continuous *b*-ion coverage with first position succinyl-K (yellow shading), and flanking *y*- and *b*-ions at the TMT labeling site (teal shading). Retention time of the selected peaks was matched to that reported in the FragPipe search output.

Histone H4 Succinylation sites: sequence coverage

Protease-specific site coverage

Succinylation sites identified by both Arg-C Ultra and r-Chymotrypsin

*r-Chymotrypsin + TMT*

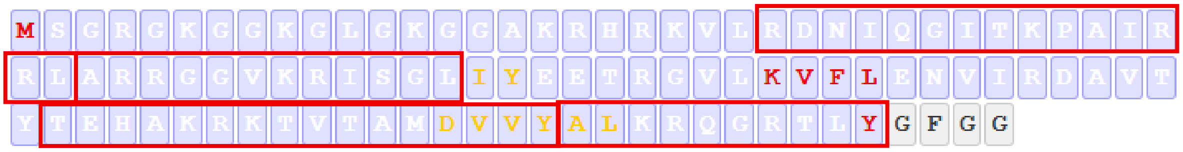


RDNIQGITkPAIRRL
ARRGGVkRISGL

TEHAKRkTVTAM;TEHAKRkTVTAMDVVY

ALkRQGRTLY; kRQGRTLY

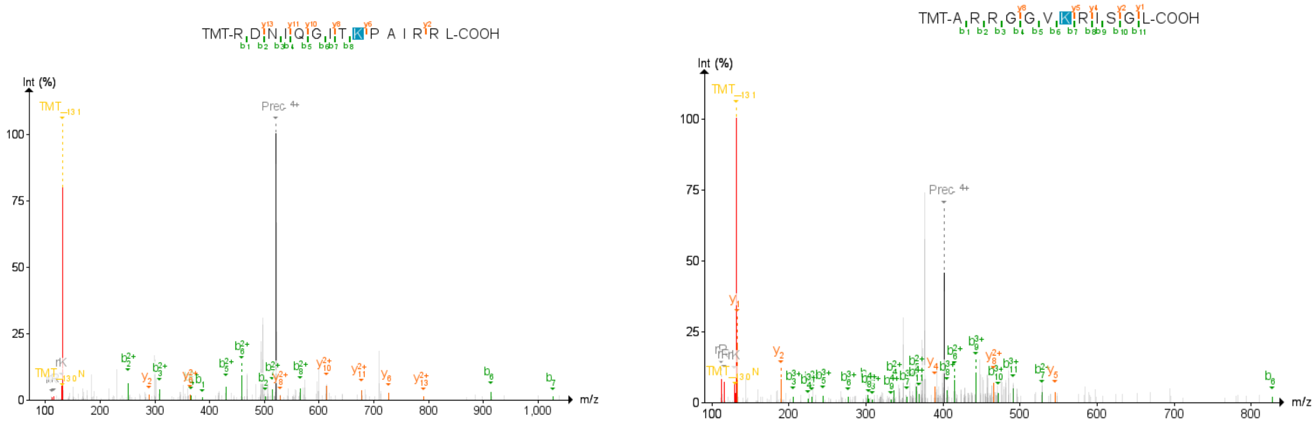


**Figure S5A.** H4 sequence coverage and succinylation sites in r-Chymotrypsin + TMT samples. Top panels: Theoretical sequence coverage is indicated by purple shading. Red boxes indicate observed peptides/sequence coverage of succinyl-K peptides. Peptides highlighted in yellow indicate protease-specific K-succinyl site coverage. Green highlighted K residues indicate succinyl-K sites identified by both r-Chymotrypsin + TMT and Arg-C Ultra + TMT. Bottom panels: Example fragment spectra of RDNIQGITkPAIRRL (+4 ion) and ARRGGVkRISGL (+4 ion) peptides showing increased *b*-ion series and flanking fragment ions of succinyl-K sites. Succinyl-K sites are highlighted in color.

*Arg-C Ultra + TMT*

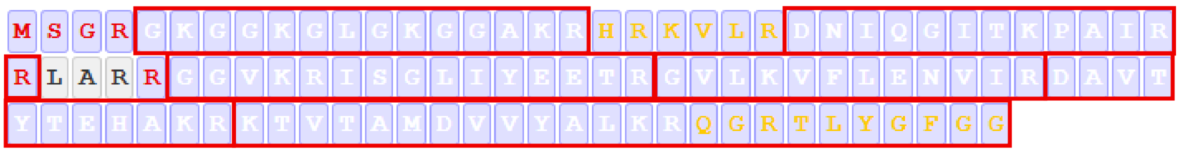


DNIQGITkPAIR; DNIQGITkPAIRR; KVLRDNIQGITkPAIR

GkGGKGLGKGGAKR

GKGGkGLGKGGAKR

DAVTYTEHAkR

kTVTAMDVVYALKR; kTVTAMDVVYALKRQGR; kTVTAMDVVYALKRQGRTLYGFGG

KTVTAMDVVYALkR

**Figure S5B.** H4 sequence coverage and succinylation sites in Arg-C Ultra + TMT samples. Top panels: Theoretical sequence coverage is indicated by purple shading. Red boxes indicate observed peptides/sequence coverage of succinyl-K peptides. Peptides highlighted in yellow indicate protease-specific K-succinyl site coverage. Green highlighted K residues indicate succinyl-K sites identified by both r-Chymotrypsin + TMT and Arg-C Ultra + TMT.

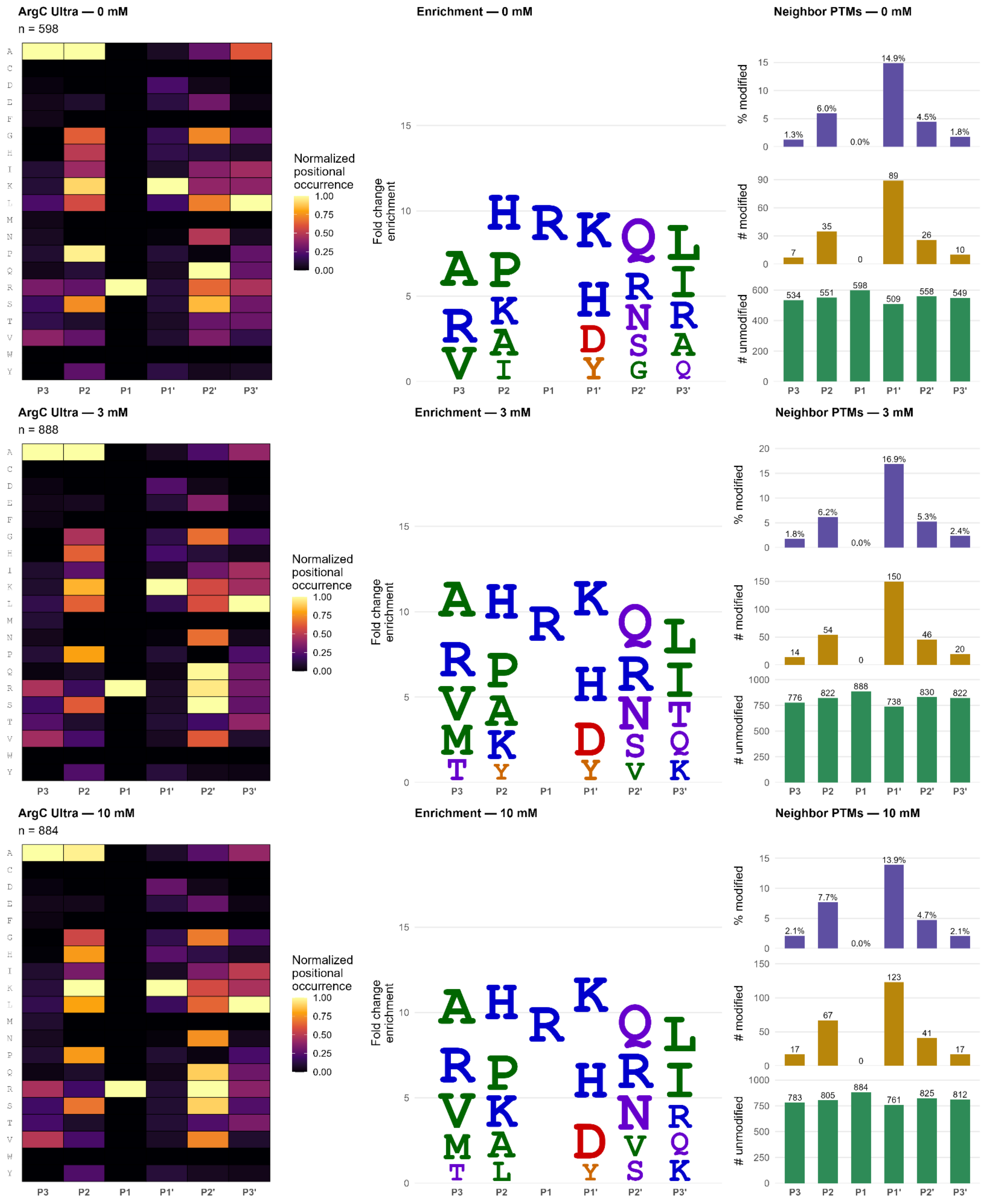


**Figure S6.** Arg-C Ultra missed cleavage motif analysis from NAM-treated and untreated HEK293T histones.

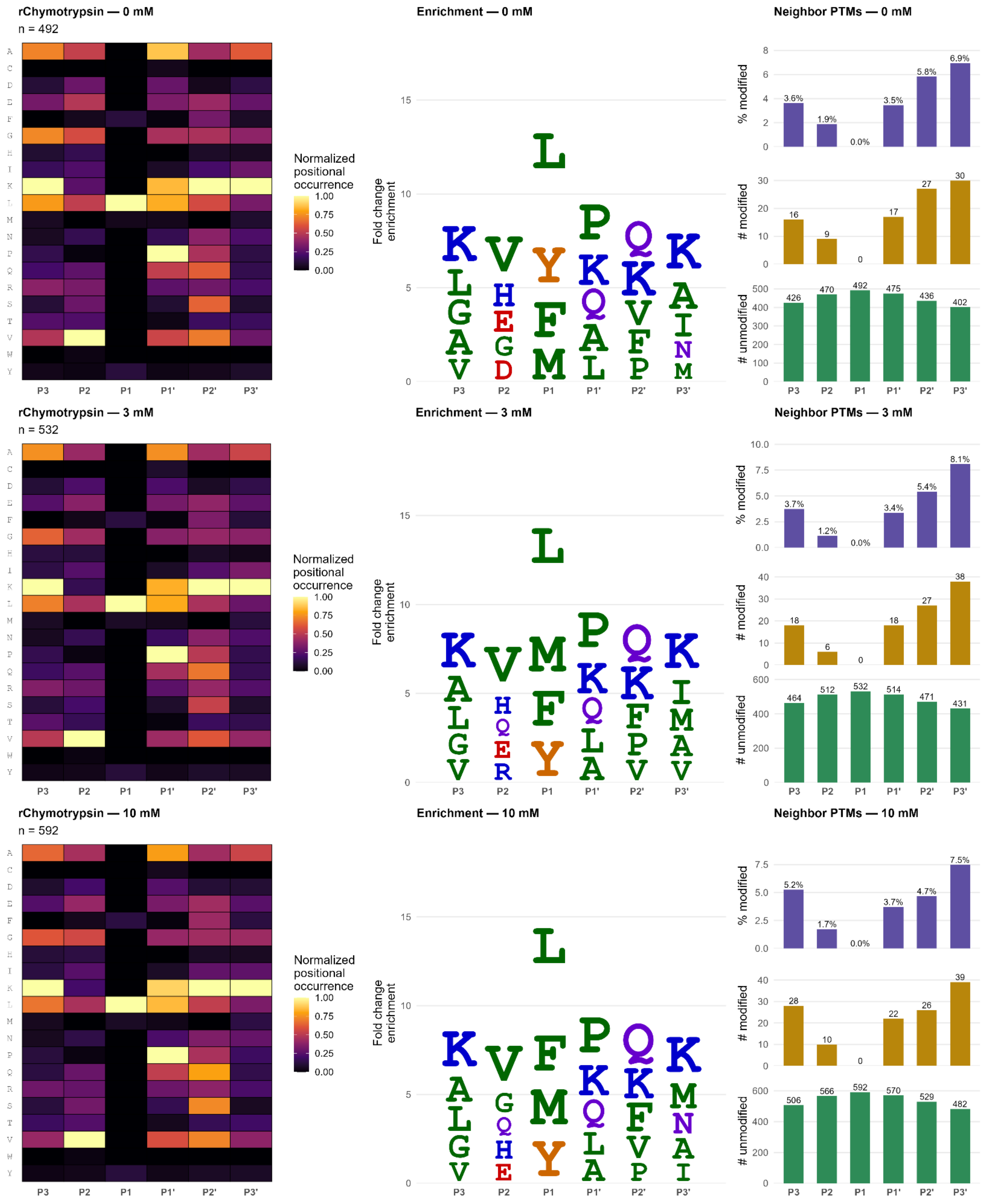


**Figure S7.** r-Chymotrypsin missed cleavage motif analysis from NAM-treated and untreated HEK293T histones.

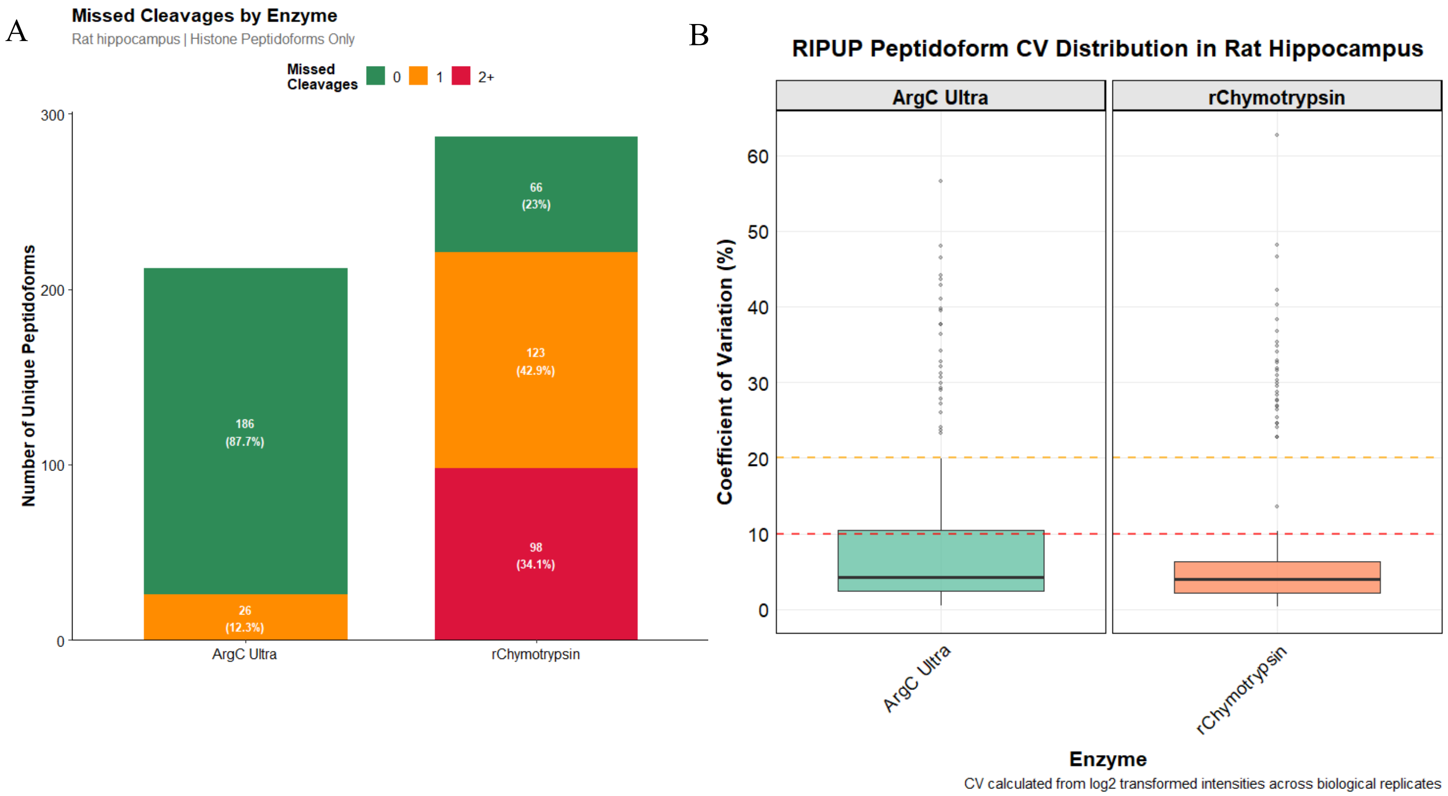


**Figure S8.** Peptidoforms, missed cleavages, and CV distributions of rat hippocampal histone peptides obtained with RIPUP. (A) Unique peptidoforms and missed cleavages in histone extracts from rat hippocampus digested with Arg-C Ultra (1:10; left) and r-Chymotrypsin (1:10; right). (B) CV distributions of peptidoforms identified with the RIPUP protocol in rat hippocampal sections.

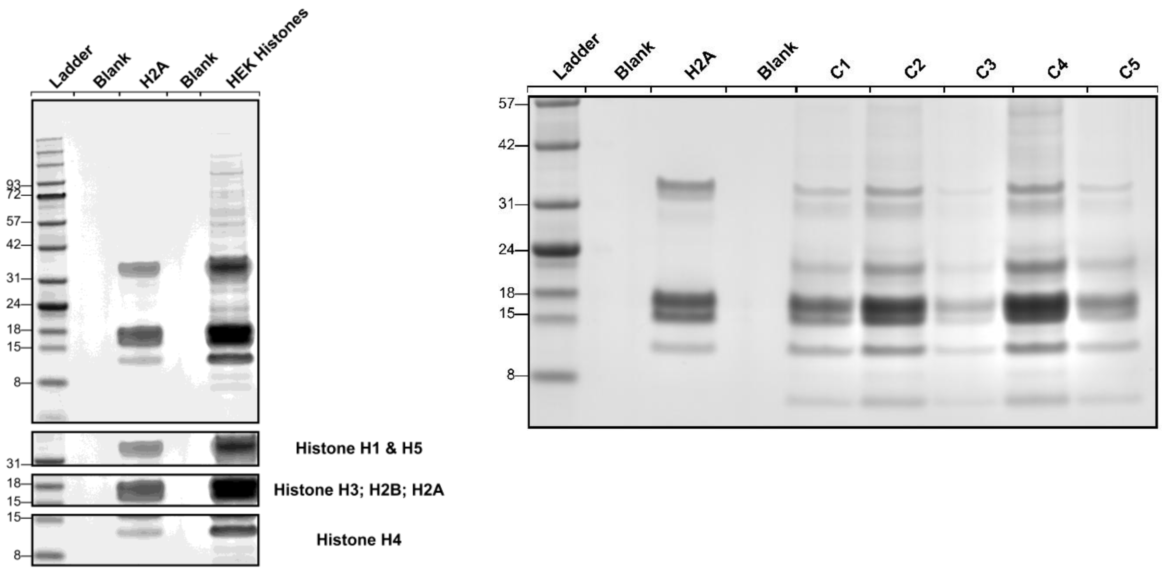


**Figure S9.** SDS-PAGE protein profiles of histones extracted from HEK293T cells and rat hippocampal sections. Five µL of extracted histones or 2 µg of histone standard were resolved on a 4-12% Bis-Tris mini-protein gel and subsequently stained to visualize histone protein bands and confirm sample integrity. Blank = no sample; C1-5 = histones extracted from rat hippocampal sections, biological replicates 1 – 5; H2A = Histone type IIA from calf thymus (catalog number H9250, Sigma-Aldrich); HEK Histones = histones extracted from HEK293T nuclei.

**Table S1:** FragPipe Detailed Mass Offsets*

| **Arg-C Ultra (unlabeled)** | | **r-Chymotrypsin (unlabeled)** | |
|---|---|---|---|
| 0 | | # Mass | Allowed sites |
| 0.984 | R | 0 | |
| 27.9949 | KST | 27.9949 | KST |
| 68.0262 | K | 68.0262 | K |
| 72.0211 | K | 72.0211 | K |
| 104.0262 | K | 104.0262 | K |
| 86.0004 | K | 86.0004 | K |
| 86.0368 | K | 86.0368 | K |
| 100.016 | K | 100.016 | K |
| 114.0317 | K | 114.0317 | K |
| 56.0262 | K | 56.0262 | K |
| 70.0419 | K | 70.0419 | K |
| 114.0429 | K | 270.1441 | K |
| 159.0684 | Q | 159.0684 | Q |
| 79.9663 | STY | 79.9663 | STY |
| 162.0528 | K | 162.0528 | K |
| 108.0211 | K | 108.0211 | K |
| 88.016 | K | 88.016 | K |
| 39.9949 | R | 39.9949 | R |
| 54.0106 | R | 54.0106 | R |
| 80.0262 | R | 80.0262 | R |
| 144.0423 | R | 144.0423 | R |
| 132.0575 | K | 132.0575 | K |

* Detailed mass offsets were adjusted according to labeling strategy as described above in 'FragPipe Settings'. Highlighted row refers to the GG (Arg-C Ultra) or RGG (r-Chymotrypsin) ubiquitin remnant from the different protease cleavage specificity, resulting in distinct delta mass shifts on K.

**Table S2:** Protease digestion conditions for histones extracted from HEK293T cells.

| Sample type | Protease | Denaturation | Derivatization (propionylation) | Digestion conditions: time (temperature) |
|---|---|---|---|---|
| HEK293 | Trypsin (control)<br>1:10 | +/- | +<br>(before and after digestion) | 6 h<br>(37 °C) |
| | Arg-C ultra<br>1:100 | +/- | +/-<br>(after digestion) | 2 h<br>(37 °C) |
| | r-Chymotrypsin<br>1:40 | +/- | +/-<br>(after digestion) | 2 h<br>(RT) |

**Table S3:** Denaturing conditions for histones extracted from HEK293T cells.*

| Enzyme | Cleavage Specificity | Denaturant | Denaturant concentration |
|---|---|---|---|
| Arg-C Ultra | R; C-term (Not before P) | Urea | 6 M (with 10 mM TCEP) |
| r-Chymotrypsin | Y, F, L, M; C-term (Not before P) | Urea | 6 M (with 5 mM TCEP) |
| Trypsin | R, K (non-propionylated); C-term | Urea | 2 M (with 5 mM TCEP) |
| Arg-C Ultra | | None | (with 10 mM TCEP) |
| r-Chymotrypsin | | | (with 5 mM TCEP) |
| Trypsin | | | (with 5 mM TCEP) |

* Trypsin is reported to retain >80% activity at 2 M urea but loses activity sharply at concentrations above 4 M (Klammer & MacCoss, 2006); accordingly, 2 M urea was used for trypsin digestions under denaturing conditions. The manufacturer's protocol for Arg-C Ultra permits digestion in buffers containing up to 6 M urea, consistent with published reports of Arg-C retaining substantial activity at 4-6 M urea (Promega Technical Manual # VA1831 and VA1832). For r-Chymotrypsin, 6 M urea was found to retain cleavage specificity and efficiency at this concentration (Adoni et al., 2026). Precise protease activity measurements as a function of urea concentration were not performed in this study. Similarly, TCEP concentrations were chosen according to the manufacturer recommendations for protease tolerances; Arg-C Ultra requires a minimum of 10 mM TCEP to keep the enzyme activated, whereas r-Chymotrypsin is a serine protease with no active-site cysteine and is therefore insensitive to the redox state of cysteines. Trypsin is similarly a serine protease for which 5 mM TCEP is sufficient for substrate disulfide reduction without adverse enzyme effects.

**Table S4:** Peptide lists of modified peptides identified in rat hippocampal sections using the RIPUP workflow (Arg-C Ultra and r-Chymotrypsin)

| Dataset_Source | Protein | Modified Sequence | Start | End | n_replicates |
|---|---|---|---|---|---|
| ArgC Ultra | H10_RAT | AKAAKKSTDHPKYSDMIVAAIQAEKNR | 15 | 41 | 4 |
| ArgC Ultra | H10_RAT | LVTTGVLKQTKGVGASGSFR | 74 | 93 | 4 |
| ArgC Ultra | H10_RAT | QSIQKYIKSHYKVGENADSQIKLSIKR | 47 | 73 | 4 |
| ArgC Ultra | H10_RAT | TENSTSTPAAKPKR | 1 | 14 | 4 |
| ArgC Ultra | H10_RAT | n[42.0106]TENSTSTPAAKPKR | 1 | 14 | 4 |
| ArgC Ultra | H11_RAT | KKPAGPSVSELIVQAVSSSKER | 34 | 55 | 4 |
| ArgC Ultra | H11_RAT | SGVSLAALKKSLAAAGYDVEKNNSR | 56 | 80 | 4 |
| ArgC Ultra | H11_RAT | SGVSLAALK[28.0313]K[28.0313]SLAAAGYDVEK[132.0575]NNSR[10.0083] | 56 | 80 | 3 |
| ArgC Ultra | H11_RAT | n[42.0106]SETAPVPQPASVAPEKPAATKKTR | 1 | 24 | 4 |
| ArgC Ultra | H14_RAT | AVKPKAAKPKTSKPKAAKPKKTAAKKK | 192 | 218 | 4 |
| ArgC Ultra | H14_RAT | KASGPPVSELITKAVAASKER | 33 | 53 | 4 |
| ArgC Ultra | H14_RAT | KASGPPVSELITK[42.0470]AVAASK[42.0470]ER[144.0423] | 33 | 53 | 3 |
| ArgC Ultra | H14_RAT | KAS[79.9663]GPPVSELITKAVAASKER | 33 | 53 | 4 |
| ArgC Ultra | H14_RAT | K[42.0106]AS[27.9949]GPPVSELITKAVAASKER | 33 | 53 | 4 |
| ArgC Ultra | H14_RAT | SETAPAAPAAPAPAEKTPIKKKAR | 1 | 24 | 4 |
| ArgC Ultra | H14_RAT | SGVSLAALKKALAAAGYDVEKNNSR | 54 | 78 | 4 |
| ArgC Ultra | H14_RAT | n[42.0106]SETAPAAPAAPAPAEKTPIKKKAR | 1 | 24 | 4 |
| ArgC Ultra | H14_RAT | n[42.0106]SETAPAAPAAPAPAEKTPIK[14.0156]KKAR | 1 | 24 | 4 |
| ArgC Ultra | H14_RAT | n[42.0106]SETAPAAPAAPAPAEKT[79.9663]PIKKKAR | 1 | 24 | 4 |
| ArgC Ultra | H14_RAT | n[42.0106]SETAPAAPAAPAPAEK[14.0156]TPIKKKAR | 1 | 24 | 3 |
| ArgC Ultra | H14_RAT | n[42.0106]SETAPAAPAAPAPAEK[42.0106]TPIKKKAR | 1 | 24 | 4 |
| ArgC Ultra | H14_RAT | n[42.0106]SET[79.9663]APAAPAAPAPAEKTPIKKKAR | 1 | 24 | 4 |
| ArgC Ultra | H15_RAT | GGVSLPALKKALAAGGYDVEKNNSR | 53 | 77 | 4 |
| ArgC Ultra | H15_RAT | KATGPPVSELITKAVSASKER | 32 | 52 | 3 |
| ArgC Ultra | H15_RAT | PKAVKSKASKPKVTKPKAAKPKAAKVKKAVSKKK | 188 | 221 | 4 |
| ArgC Ultra | H15_RAT | SETAPAETTAPAPVEKSPAKKKTKKAGAAKR | 1 | 31 | 4 |
| ArgC Ultra | H2A1C_RAT | AGLQFPVGR | 21 | 29 | 4 |
| ArgC Ultra | H2A1C_RAT | HLQLAIR | 82 | 88 | 4 |
| ArgC Ultra | H2A1C_RAT | NDEELNKLLGR | 89 | 99 | 4 |

| | | | | | |
|---|---|---|---|---|---|
| ArgC Ultra | H2A1C_RAT | NDEELNK[27.9949]LLGR | 89 | 99 | 3 |
| ArgC Ultra | H2A1C_RAT | VGAGAPVYLAAVLEYLTAEILELAGNAAR | 43 | 71 | 3 |
| ArgC Ultra | H2A1C_RAT | VTIAQGGVLPNIQAVLLPKKTESHHKAKGK | 100 | 129 | 4 |
| ArgC Ultra | H2A1C_RAT | VTIAQGGVLPNIQAVLLPKKTESHHKAKGK[42.0106] | 100 | 129 | 4 |
| ArgC Ultra | H2A1C_RAT | VTIAQGGVLPNIQAVLLPKKTESHHKAK[42.0106]GK | 100 | 129 | 4 |
| ArgC Ultra | H2A1C_RAT | VTIAQGGVLPNIQAVLLPKKTESHHK[42.0106]AKGK | 100 | 129 | 3 |
| ArgC Ultra | H2A1C_RAT | VTIAQGGVLPNIQAVLLPKK[114.0429]TESHHKAKGK | 100 | 129 | 4 |
| ArgC Ultra | H2A1C_RAT | VTIAQGGVLPNIQAVLLPK[114.0429]KTESHHKAKGK | 100 | 129 | 3 |
| ArgC Ultra | H2A1C_RAT | VTIAQGGVLPNIQAVLLPK[14.0156]K[72.0211]TESHHK[28.0313]AKGK | 100 | 129 | 4 |
| ArgC Ultra | H2A1C_RAT | VTIAQGGVLPNIQAVLLPK[42.0106]KTESHHKAKGK | 100 | 129 | 3 |
| ArgC Ultra | H2A1C_RAT | VTIAQGGVLPNIQAVLLPK[42.0470]K[72.0211]TESHHKAKGK | 100 | 129 | 4 |
| ArgC Ultra | H2A1C_RAT | VTIAQGGVLPNIQ[159.0684]AVLLPK[28.0313]K[28.0313]TESHHKAKGK | 100 | 129 | 4 |
| ArgC Ultra | H2A1C_RAT | VTIAQGGVLPNIQ[159.0684]AVLLPK[42.0470]K[14.0156]TESHHKAKGK | 100 | 129 | 3 |
| ArgC Ultra | H2A1F_RAT | VTIAQGGVLPNIQAVLLPKKTESHHKPKGK | 100 | 129 | 4 |
| ArgC Ultra | H2A1F_RAT | VTIAQGGVLPNIQAVLLPKKTESHHKPK[42.0106]GK | 100 | 129 | 4 |
| ArgC Ultra | H2A1F_RAT | VTIAQGGVLPNIQAVLLPKKTESHHK[114.0429]PKGK | 100 | 129 | 4 |
| ArgC Ultra | H2A1F_RAT | VTIAQGGVLPNIQAVLLPK[42.0470]K[104.0262]TESHHK[42.0106]PKGK | 100 | 129 | 3 |
| ArgC Ultra | H2A1F_RAT | VTIAQGGVLPNIQAVLLPK[88.0160]KTESHHKPKGK | 100 | 129 | 2 |
| ArgC Ultra | H2A2A_RAT | NDEELNKLLGKVTIAQGGVLPNIQAVLLPKKTESHHKAKGK | 89 | 129 | 4 |
| ArgC Ultra | H2A2A_RAT | NDEELNKLLGKVTIAQGGVLPNIQAVLLPKKTESHHKAKGK[42.0106] | 89 | 129 | 4 |
| ArgC Ultra | H2A2A_RAT | NDEELNKLLGKVTIAQGGVLPNIQAVLLPKKTESHHKAK[42.0106]GK | 89 | 129 | 4 |

| | | | | | |
|---|---|---|---|---|---|
| ArgC Ultra | H2A2A_RAT | NDEELNKLLGKVTIAQGGVLPNIQAVLLPK[42.0470]K[72.0211]TESHHKAKGK | 89 | 129 | 4 |
| ArgC Ultra | H2A2A_RAT | NDEELNKLLGKVTIAQGGVLPNIQAVLLPK[72.0211]KTESHHK[42.0106]AKGK | 89 | 129 | 2 |
| ArgC Ultra | H2A2A_RAT | NDEELNKLLGKVTIAQGGVLPNIQAVLLPK[72.0211]K[14.0156]TESHHK[14.0156]AK[14.0156]GK | 89 | 129 | 3 |
| ArgC Ultra | H2A2A_RAT | VGAGAPVYMAAVLEYLTAEILELAGNAAR | 43 | 71 | 3 |
| ArgC Ultra | H2A4_RAT | VTIAQGGVLPNIQAVLLPKKTESHHKSQTK | 100 | 129 | 4 |
| ArgC Ultra | H2A4_RAT | VTIAQGGVLPNIQAVLLPK[28.0313]K[68.0262]TESHHKSQTK | 100 | 129 | 3 |
| ArgC Ultra | H2AJ_RAT | VTIAQGGVLPNIQAVLLPKKTESQKVKSK | 100 | 128 | 4 |
| ArgC Ultra | H2AJ_RAT | VTIAQGGVLPNIQAVLLPKKTESQK[42.0470]VK[88.0160]SK | 100 | 128 | 3 |
| ArgC Ultra | H2AJ_RAT | VTIAQGGVLPNIQAVLLPK[114.0429]KTESQKVKSK | 100 | 128 | 3 |
| ArgC Ultra | H2AJ_RAT | VTIAQGGVLPNIQAVLLPK[14.0156]K[42.0106]TESQK[68.0262]VK[42.0106]SK[28.0313] | 100 | 128 | 3 |
| ArgC Ultra | H2AJ_RAT | VTIAQGGVLPNIQAVLLPK[42.0106]KTESQKVKSK | 100 | 128 | 4 |
| ArgC Ultra | H2AJ_RAT | VTIAQGGVLPNIQAVLLPK[42.0106]K[42.0106]TESQK[14.0156]VK[104.0262]SK | 100 | 128 | 4 |
| ArgC Ultra | H2AJ_RAT | VTIAQGGVLPNIQAVLLPK[42.0470]K[72.0211]TESQKVKSK | 100 | 128 | 4 |
| ArgC Ultra | H2AJ_RAT | VTIAQGGVLPNIQAVLLPK[88.0160]KTESQKVKSK | 100 | 128 | 4 |
| ArgC Ultra | H2AY_RAT | KLKSIAFPSIGSGR | 300 | 313 | 4 |
| ArgC Ultra | H2AY_RAT | SAKAGVIFPVGR | 15 | 26 | 4 |
| ArgC Ultra | H2AZ_RAT | AGGKAGK[42.0106]DSGKAKTKAVSR | 1 | 19 | 3 |
| ArgC Ultra | H2AZ_RAT | AGGKAGK[42.0106]DSGK[42.0106]AKTKAVSR | 1 | 19 | 4 |
| ArgC Ultra | H2AZ_RAT | AGGK[42.0106]AGK[42.0106]DSGKAKTKAVSR | 1 | 19 | 4 |
| ArgC Ultra | H2AZ_RAT | AGGK[42.0106]AGK[42.0106]DSGK[42.0106]AKTKAVSR | 1 | 19 | 4 |
| ArgC Ultra | H2AZ_RAT | GDEELDSLIKATIAGGGVIPHIHKSLIGKKGQQKTV | 92 | 127 | 4 |
| ArgC Ultra | H2AZ_RAT | GDEELDSLIKATIAGGGVIPHIHKSLIGK[42.0470]K[72.0211]GQQKTV | 92 | 127 | 4 |
| ArgC Ultra | H2AZ_RAT | GDEELDSLIKATIAGGGVIPHIHK[42.0470]SLIGK[72.0211]KGQQKTV | 92 | 127 | 3 |
| ArgC Ultra | H2AZ_RAT | GDEELDSLIKATIAGGGVIPHIHK[72.0211]SLIGK[42.0106]KGQQKTV | 92 | 127 | 2 |
| ArgC Ultra | H2B1_RAT | EIQTAVR | 92 | 98 | 4 |
| ArgC Ultra | H2B1_RAT | KESYSVYVYKVLKQVHPDTGISSKAMGIMNSFVNDIFER | 34 | 72 | 3 |
| ArgC Ultra | H2B1_RAT | K[14.0156]ESYSVYVYKVLKQVHPDTGISSKAMGIMNSFVNDIFER | 34 | 72 | 3 |
| ArgC Ultra | H2B1_RAT | LLLPGELAKHAVSEGTKAVTKYTSSK | 99 | 124 | 4 |
| ArgC Ultra | H2B1_RAT | LLLPGELAKHAVSEGTKAVTKYTSSK[114.0429] | 99 | 124 | 4 |
| ArgC Ultra | H2B1_RAT | LLLPGELAKHAVSEGTKAVTK[114.0429]YTSSK | 99 | 124 | 4 |

| | | | | | |
|---|---|---|---|---|---|
| ArgC Ultra | H2B1_RAT | LLLPGELAKHAVSEGTKAVTK[42.0106]YTSSK[72.0211] | 99 | 124 | 3 |
| ArgC Ultra | H2B1_RAT | LLLPGELAK[27.9949]HAVSEGTKAVTKYTSSK | 99 | 124 | 4 |
| ArgC Ultra | H2B1_RAT | PEPAKSR[39.9949]PAPKKGSKKAVTKAQKKDGKER | 1 | 29 | 3 |
| ArgC Ultra | H31_RAT | K[28.0313]SAPATGGVKKPHR | 27 | 40 | 4 |
| ArgC Ultra | H31_RAT | K[28.0313]SAPATGGVK[14.0156]KPHR | 27 | 40 | 4 |
| ArgC Ultra | H31_RAT | K[28.0313]SAPATGGVK[28.0313]KPHR | 27 | 40 | 3 |
| ArgC Ultra | H31_RAT | K[42.0470]SAPATGGVKKPHR | 27 | 40 | 4 |
| ArgC Ultra | H31_RAT | K[42.0470]SAPATGGVK[28.0313]KPHR | 27 | 40 | 4 |
| ArgC Ultra | H33_RAT | EIAQDFKTDLR | 73 | 83 | 4 |
| ArgC Ultra | H33_RAT | EIAQDFK[14.0156]TDLR | 73 | 83 | 4 |
| ArgC Ultra | H33_RAT | EIAQDFK[28.0313]TDLR | 73 | 83 | 4 |
| ArgC Ultra | H33_RAT | KLPFQR | 64 | 69 | 4 |
| ArgC Ultra | H33_RAT | KQLATK[42.0106]AAR | 18 | 26 | 4 |
| ArgC Ultra | H33_RAT | KSAPSTGGVKKPHR | 27 | 40 | 4 |
| ArgC Ultra | H33_RAT | KSAPSTGGVK[28.0313]KPHR | 27 | 40 | 4 |
| ArgC Ultra | H33_RAT | KSAPSTGGVK[42.0470]KPHR | 27 | 40 | 4 |
| ArgC Ultra | H33_RAT | K[14.0156]S[27.9949]APSTGGVK[14.0156]KPHR | 27 | 40 | 4 |
| ArgC Ultra | H33_RAT | K[14.0156]S[27.9949]APSTGGVK[28.0313]KPHR | 27 | 40 | 4 |
| ArgC Ultra | H33_RAT | K[14.0156]S[27.9949]APSTGGVK[42.0470]KPHR | 27 | 40 | 3 |
| ArgC Ultra | H33_RAT | K[28.0313]SAPSTGGVK[14.0156]KPHR | 27 | 40 | 4 |
| ArgC Ultra | H33_RAT | K[28.0313]SAPSTGGVK[28.0313]KPHR | 27 | 40 | 4 |
| ArgC Ultra | H33_RAT | K[42.0106]QLATKAAR | 18 | 26 | 4 |
| ArgC Ultra | H33_RAT | K[42.0106]QLATK[42.0106]AAR | 18 | 26 | 4 |
| ArgC Ultra | H33_RAT | K[42.0106]SAPSTGGVK[28.0313]KPHR | 27 | 40 | 3 |
| ArgC Ultra | H33_RAT | K[42.0106]STGGK[42.0106]APR | 9 | 17 | 4 |
| ArgC Ultra | H33_RAT | K[42.0470]SAPSTGGVK[14.0156]KPHR | 27 | 40 | 4 |
| ArgC Ultra | H33_RAT | K[42.0470]SAPSTGGVK[28.0313]KPHR | 27 | 40 | 4 |
| ArgC Ultra | H33_RAT | VTIMPKDIQLAR | 117 | 128 | 4 |
| ArgC Ultra | H33_RAT | YQKSTELLIR | 54 | 63 | 4 |
| ArgC Ultra | H4_RAT | DAVTYTEHAKR | 68 | 78 | 4 |
| ArgC Ultra | H4_RAT | DNIQGITKPAIR | 24 | 35 | 4 |
| ArgC Ultra | H4_RAT | GKGGKGLGK[42.0106]GGAK[42.0106]R | 4 | 17 | 4 |

| | | | | | |
|---|---|---|---|---|---|
| ArgC Ultra | H4_RAT | GKGGK[42.0106]GLGKGGAK[42.0106]R | 4 | 17 | 4 |
| ArgC Ultra | H4_RAT | GKGGK[42.0106]GLGK[42.0106]GGAKR | 4 | 17 | 4 |
| ArgC Ultra | H4_RAT | GKGGK[42.0106]GLGK[42.0106]GGAK[42.0106]R | 4 | 17 | 4 |
| ArgC Ultra | H4_RAT | GK[42.0106]GGKGLGKGGAK[42.0106]R | 4 | 17 | 4 |
| ArgC Ultra | H4_RAT | GK[42.0106]GGKGLGK[42.0106]GGAK[42.0106]R | 4 | 17 | 4 |
| ArgC Ultra | H4_RAT | GK[42.0106]GGK[42.0106]GLGKGGAK[42.0106]R | 4 | 17 | 4 |
| ArgC Ultra | H4_RAT | GK[42.0106]GGK[42.0106]GLGK[42.0106]GGAK[27.9949]R[14.0156] | 4 | 17 | 4 |
| ArgC Ultra | H4_RAT | GK[42.0106]GGK[42.0106]GLGK[42.0106]GGAK[42.0106]R | 4 | 17 | 4 |
| ArgC Ultra | H4_RAT | GVLKVFLENVIR | 56 | 67 | 4 |
| ArgC Ultra | H4_RAT | GVLK[162.0528]VFLENVIR | 56 | 67 | 4 |
| ArgC Ultra | H4_RAT | ISGLIYEETR | 46 | 55 | 4 |
| ArgC Ultra | H4_RAT | IS[27.9949]GLIYEETR | 46 | 55 | 4 |
| ArgC Ultra | H4_RAT | IS[79.9663]GLIYEETR | 46 | 55 | 4 |
| ArgC Ultra | H4_RAT | KTVTAMDVVYALKR | 79 | 92 | 4 |
| ArgC Ultra | H4_RAT | KTVTAMDVVYALK[162.0528]R | 79 | 92 | 4 |
| ArgC Ultra | H4_RAT | KT[27.9949]VTAMDVVYALKR | 79 | 92 | 4 |
| ArgC Ultra | H4_RAT | K[162.0528]TVTAMDVVYALKR | 79 | 92 | 4 |
| ArgC Ultra | H4_RAT | K[42.0106]T[27.9949]VTAMDVVYALKR | 79 | 92 | 4 |
| ArgC Ultra | H4_RAT | SGR[14.0156]GK[42.0106]GGKGLGKGGAK[42.0106]R | 1 | 17 | 4 |
| ArgC Ultra | H4_RAT | n[42.0106]SGR[14.0156]GKGGKGLGKGGAK[42.0106]R | 1 | 17 | 4 |
| ArgC Ultra | H4_RAT | n[42.0106]SGR[14.0156]GKGGK[42.0106]GLGKGGAK[42.0106]R | 1 | 17 | 4 |
| ArgC Ultra | H4_RAT | n[42.0106]SGR[14.0156]GK[14.0156]GGKGLGKGGAK[42.0106]R | 1 | 17 | 4 |
| ArgC Ultra | H4_RAT | n[42.0106]SGR[28.0313]GKGGKGLGKGGAK[42.0106]R | 1 | 17 | 4 |
| rChymotrypsin | H10_RAT | AKGDEPKRSVAF | 95 | 106 | 4 |
| rChymotrypsin | H10_RAT | KQTKGVGASGSF | 81 | 92 | 4 |
| rChymotrypsin | H10_RAT | KQTKGVGASGS[-18.0106]F | 81 | 92 | 4 |
| rChymotrypsin | H10_RAT | KVGENADSQIKL | 58 | 69 | 4 |
| rChymotrypsin | H10_RAT | KVGENADSQIKLSIKRLVTTGVL | 58 | 80 | 4 |
| rChymotrypsin | H10_RAT | KVGENADS[-18.0106]QIKL | 58 | 69 | 4 |
| rChymotrypsin | H10_RAT | RLAKGDEPKRSVAF | 93 | 106 | 4 |
| rChymotrypsin | H10_RAT | SDMIVAAIQAEKNRAGSSRQSIQKY | 28 | 52 | 4 |
| rChymotrypsin | H10_RAT | SDM[15.9949]IVAAIQAEKNRAGSSRQSIQKY | 28 | 52 | 4 |

| | | | | | |
|---|---|---|---|---|---|
| rChymotrypsin | H10_RAT | SDM[15.9949]IVAAIQAEK[14.0156]NR[39.9949]AGS[-18.0106]S[-18.0106]RQS[-18.0106]IQKY | 28 | 52 | 2 |
| rChymotrypsin | H10_RAT | SDM[15.9949]IVAAIQAEK[42.0106]NR[10.0083]AGS[-18.0106]S[-18.0106]RQSIQKY | 28 | 52 | 3 |
| rChymotrypsin | H10_RAT | SIKRLVTTGVL | 70 | 80 | 4 |
| rChymotrypsin | H10_RAT | SIKRLVTTGVLKQTKGVGASGSF | 70 | 92 | 4 |
| rChymotrypsin | H10_RAT | TENSTSTPAAKPKRAKAAKKSTDHPKY | 1 | 27 | 4 |
| rChymotrypsin | H10_RAT | VTTGVLKQTKGVGASGSF | 75 | 92 | 4 |
| rChymotrypsin | H11_RAT | KK[42.0106]SLAAAGY | 64 | 72 | 3 |
| rChymotrypsin | H11_RAT | KSLVNKGTLVQTKGTGAAGSF | 86 | 106 | 4 |
| rChymotrypsin | H14_RAT | AAAGYDVEKNNSRIKL | 66 | 81 | 4 |
| rChymotrypsin | H14_RAT | AAAGYDVEKNNSRIKLGL | 66 | 83 | 4 |
| rChymotrypsin | H14_RAT | AAAGYDVEK[114.0317]NNSR[28.0313]IK[28.0313]L | 66 | 81 | 3 |
| rChymotrypsin | H14_RAT | AALKKALAAAGY | 59 | 70 | 4 |
| rChymotrypsin | H14_RAT | DVEKNNSRIKL | 71 | 81 | 4 |
| rChymotrypsin | H14_RAT | DVEKNNSRIKLGL | 71 | 83 | 4 |
| rChymotrypsin | H14_RAT | DVEKNNSRIKLGLKSL | 71 | 86 | 4 |
| rChymotrypsin | H14_RAT | GLKSLVSKGTL | 82 | 92 | 4 |
| rChymotrypsin | H14_RAT | ITKAVAASKERSGVSL | 43 | 58 | 4 |
| rChymotrypsin | H14_RAT | ITKAVAASKERSGVSLAAL | 43 | 61 | 4 |
| rChymotrypsin | H14_RAT | ITKAVAASKERSGVSLAALKKAL | 43 | 65 | 4 |
| rChymotrypsin | H14_RAT | KKALAAAGY | 62 | 70 | 4 |
| rChymotrypsin | H14_RAT | KKALAAAGYDVEKNNSRIKL | 62 | 81 | 4 |
| rChymotrypsin | H14_RAT | KKALAAAGYDVEK[114.0317]NNSR[14.0156]IK[42.0470]L | 62 | 81 | 2 |
| rChymotrypsin | H14_RAT | KKALAAAGYDVEK[132.0575]NNSR[10.0083]IK[28.0313]L | 62 | 81 | 4 |
| rChymotrypsin | H14_RAT | KKALAAAGYDVEK[70.0419]NNSR[28.0313]IK[72.0211]L | 62 | 81 | 4 |
| rChymotrypsin | H14_RAT | KSLVSKGTL | 84 | 92 | 4 |
| rChymotrypsin | H14_RAT | KSLVSKGTLVQTKGTGASGSF | 84 | 104 | 4 |
| rChymotrypsin | H14_RAT | KSLVSKGTLVQTKGTGASGS[-18.0106]F | 84 | 104 | 3 |
| rChymotrypsin | H14_RAT | K[14.0156]SLVSK[114.0317]GTLVQTK[42.0470]GTGASGSF | 84 | 104 | 2 |
| rChymotrypsin | H14_RAT | K[14.0156]SLVSK[70.0419]GTLVQTK[86.0368]GTGASGSF | 84 | 104 | 3 |
| rChymotrypsin | H14_RAT | K[42.0106]SLVSK[42.0470]GTLVQTK[86.0368]GTGASGSF | 84 | 104 | 4 |
| rChymotrypsin | H14_RAT | K[70.0419]S[27.9949]LVSK[42.0470]GTLVQTK[42.0470]GTGASGSF | 84 | 104 | 3 |
| rChymotrypsin | H14_RAT | VQTKGTGASGSF | 93 | 104 | 4 |

| | | | | | |
|---|---|---|---|---|---|
| rChymotrypsin | H14_RAT | VSKGTLVQTKGTGASGSF | 87 | 104 | 4 |
| rChymotrypsin | H15_RAT | AAGGYDVEKNNSRIKL | 65 | 80 | 4 |
| rChymotrypsin | H15_RAT | AAGGYDVEKNNSRIKLGL | 65 | 82 | 4 |
| rChymotrypsin | H15_RAT | ITKAVSASKERGGVSLPAL | 42 | 60 | 4 |
| rChymotrypsin | H15_RAT | ITKAVSASKERGGVSLPALKKAL | 42 | 64 | 3 |
| rChymotrypsin | H15_RAT | KKALAAGGYDVEKNNSRIKL | 61 | 80 | 4 |
| rChymotrypsin | H15_RAT | KKALAAGGYDVEK[114.0317]NNSR[14.0156]IK[42.0470]L | 61 | 80 | 2 |
| rChymotrypsin | H15_RAT | KKALAAGGYDVEK[70.0419]NNSR[28.0313]IK[72.0211]L | 61 | 80 | 4 |
| rChymotrypsin | H2A1F_RAT | LRKGNYSERVGAGAPVY | 34 | 50 | 4 |
| rChymotrypsin | H2A1F_RAT | RKGNYSERVGAGAPVY | 35 | 50 | 4 |
| rChymotrypsin | H2A1F_RAT | RKGNYSERVGAGAPVYL | 35 | 51 | 4 |
| rChymotrypsin | H2A1F_RAT | SERVGAGAPVY | 40 | 50 | 4 |
| rChymotrypsin | H2A1F_RAT | SERVGAGAPVYL | 40 | 51 | 4 |
| rChymotrypsin | H2A1F_RAT | SGRGKQGGK[27.9949]ARAKAKTRSSRAGLQFPVGRVHRL | 1 | 33 | 4 |
| rChymotrypsin | H2A1F_RAT | TAEILELAANAAR[144.0423]DNK[42.0470]K[14.0156]TR[14.0156]IIPR[14.0156]HL | 59 | 83 | 4 |
| rChymotrypsin | H2A2A_RAT | AERVGAGAPVY | 40 | 50 | 4 |
| rChymotrypsin | H2A2A_RAT | AERVGAGAPVYM | 40 | 51 | 4 |
| rChymotrypsin | H2A2A_RAT | AERVGAGAPVYM[15.9949] | 40 | 51 | 4 |
| rChymotrypsin | H2A2A_RAT | AGNAARDNKKTRIIPRHL | 66 | 83 | 4 |
| rChymotrypsin | H2A2A_RAT | AGNAARDNKKTRIIPRHLQL | 66 | 85 | 4 |
| rChymotrypsin | H2A2A_RAT | AIRNDEEL | 86 | 93 | 4 |
| rChymotrypsin | H2A2A_RAT | AIRNDEELNKL | 86 | 96 | 4 |
| rChymotrypsin | H2A2A_RAT | AIRNDEELNKLL | 86 | 97 | 4 |
| rChymotrypsin | H2A2A_RAT | ELAGNAARDNKKTRIIPRHL | 64 | 83 | 4 |

| | | | | | |
|---|---|---|---|---|---|
| rChymotrypsin | H2A2A_RAT | ELAGNAARDNKKTRIIPRHLQL | 64 | 85 | 4 |
| rChymotrypsin | H2A2A_RAT | E[-18.0106]LAGNAARDNKKTRIIPRHL | 64 | 83 | 4 |
| rChymotrypsin | H2A2A_RAT | E[-18.0106]LAGNAARDNKKTRIIPRHLQL | 64 | 85 | 4 |
| rChymotrypsin | H2A2A_RAT | LRKGNYAERVGAGAPVY | 34 | 50 | 4 |
| rChymotrypsin | H2A2A_RAT | PNIQAVLLPKKTESHHKAKGK | 109 | 129 | 4 |
| rChymotrypsin | H2A2A_RAT | PVGRVHRL | 26 | 33 | 4 |
| rChymotrypsin | H2A2A_RAT | PVGRVHRLL | 26 | 34 | 4 |
| rChymotrypsin | H2A2A_RAT | PVGRVHRLLRKGNY | 26 | 39 | 4 |
| rChymotrypsin | H2A2A_RAT | QFPVGRVHRL | 24 | 33 | 4 |
| rChymotrypsin | H2A2A_RAT | QFPVGRVHRLL | 24 | 34 | 4 |
| rChymotrypsin | H2A2A_RAT | QLAIRNDEEL | 84 | 93 | 3 |
| rChymotrypsin | H2A2A_RAT | QLAIRNDEELNKL | 84 | 96 | 4 |
| rChymotrypsin | H2A2A_RAT | RKGNYAERVGAGAPVY | 35 | 50 | 4 |
| rChymotrypsin | H2A2A_RAT | RKGNYAERVGAGAPVYM | 35 | 51 | 4 |
| rChymotrypsin | H2A2A_RAT | RKGNYAERVGAGAPVYM[15.9949] | 35 | 51 | 4 |
| rChymotrypsin | H2A2A_RAT | TAEILELAGNAARDNKKTRIIPRHL | 59 | 83 | 4 |
| rChymotrypsin | H2A2A_RAT | TAEILELAGNAAR[10.0083]DNK[70.0419]K[56.0262]TR[10.0083]IIPR[10.0083]HL | 59 | 83 | 4 |
| rChymotrypsin | H2AJ_RAT | AERVGAGAPVYL | 40 | 51 | 4 |
| rChymotrypsin | H2AJ_RAT | RKGNYAERVGAGAPVYL | 35 | 51 | 4 |
| rChymotrypsin | H2AY_RAT | AVANDEELNQL | 83 | 93 | 3 |
| rChymotrypsin | H2AY_RAT | AVANDEELNQLL | 83 | 94 | 4 |
| rChymotrypsin | H2AY_RAT | ELAGNAARDNKKGRVTPRHILL | 61 | 82 | 4 |
| rChymotrypsin | H2AY_RAT | ELAGNAAR[10.0083]DNK[28.0313]KGR[10.0083]VTPR[10.0083]HILL | 61 | 82 | 4 |
| rChymotrypsin | H2AY_RAT | ELAGNAAR[10.0083]DNK[56.0262]KGR[10.0083]VTPR[10.0083]HILL | 61 | 82 | 4 |

| | | | | | |
|---|---|---|---|---|---|
| rChymotrypsin | H2AY_RAT | ILKAISSY | 325 | 332 | 3 |
| rChymotrypsin | H2AY_RAT | KGVTIASGGVLPNIHPELL | 95 | 113 | 4 |
| rChymotrypsin | H2AY_RAT | M[15.9949]AAVLEY | 48 | 54 | 2 |
| rChymotrypsin | H2AY_RAT | RIGVGAPVY | 39 | 47 | 4 |
| rChymotrypsin | H2AY_RAT | RKKNGPLEVAGAAVSAGHGLPAKF | 244 | 267 | 4 |
| rChymotrypsin | H2AY_RAT | R[144.0423]K[28.0313]K[70.0419]NGPLEVAGAAVSAGHGLPAKF | 244 | 267 | 4 |
| rChymotrypsin | H2AY_RAT | SSSIKTVY | 338 | 345 | 3 |
| rChymotrypsin | H2AY_RAT | SSSIKTVYF | 338 | 346 | 4 |
| rChymotrypsin | H2AY_RAT | STKSLFL | 185 | 191 | 3 |
| rChymotrypsin | H2AY_RAT | TAEILELAGNAARDNKKGRVTPRHIL | 56 | 81 | 3 |
| rChymotrypsin | H2AY_RAT | TAEILELAGNAARDNK[28.0313]K[14.0156]GR[14.0156]VTPR[14.0156]HIL | 56 | 81 | 4 |
| rChymotrypsin | H2AY_RAT | TAEILELAGNAARDNK[70.0419]K[86.0368]GR[14.0156]VTPR[28.0313]HIL | 56 | 81 | 4 |
| rChymotrypsin | H2AY_RAT | TAEILELAGNAAR[10.0083]DNK[28.0313]K[70.0419]GR[10.0083]VTPR[80.0262]HIL | 56 | 81 | 4 |
| rChymotrypsin | H2AY_RAT | TAEILELAGNAAR[14.0156]DNK[42.0470]K[14.0156]GRVTPRHIL | 56 | 81 | 4 |
| rChymotrypsin | H2AY_RAT | TAEILELAGNAAR[28.0313]DNK[70.0419]K[42.0470]GRVTPR[39.9949]HIL | 56 | 81 | 4 |
| rChymotrypsin | H2AY_RAT | TVLSTKSLF | 182 | 190 | 4 |
| rChymotrypsin | H2AY_RAT | VQEMAKLDAN | 358 | 367 | 4 |
| rChymotrypsin | H2AY_RAT | VQEM[15.9949]AKLDAN | 358 | 367 | 4 |
| rChymotrypsin | H2AY_RAT | VSTM[15.9949]SSSIKTVYF | 334 | 346 | 4 |
| rChymotrypsin | H2AZ_RAT | AGGKAGKDSGKAKTKAVSRSQRAGL | 1 | 25 | 4 |
| rChymotrypsin | H2AZ_RAT | AGGKAGKDSGKAKTKAVSRSQRAGLQFPVGRIHRHL | 1 | 36 | 4 |
| rChymotrypsin | H2AZ_RAT | AGNASKDLKVKRITPRHLQL | 69 | 88 | 4 |
| rChymotrypsin | H2AZ_RAT | AGNASKDLK[28.0313]VK[70.0419]R[144.0423]ITPRHLQL | 69 | 88 | 4 |
| rChymotrypsin | H2AZ_RAT | AIRGDEELDSL | 89 | 99 | 4 |
| rChymotrypsin | H2AZ_RAT | AIRGDEELDSLIKATIAGGGVIPHIHKSL | 89 | 117 | 2 |
| rChymotrypsin | H2AZ_RAT | ELAGNASKDL | 67 | 76 | 4 |
| rChymotrypsin | H2AZ_RAT | ELAGNASKDLKVKRITPRHL | 67 | 86 | 4 |
| rChymotrypsin | H2AZ_RAT | IKATIAGGGVIPHIHKSL | 100 | 117 | 4 |
| rChymotrypsin | H2AZ_RAT | IKATIAGGGVIPHIHKSLIGKKGQQKTV | 100 | 127 | 4 |
| rChymotrypsin | H2AZ_RAT | KSRTTSHGRVGATAAVY | 37 | 53 | 4 |
| rChymotrypsin | H2AZ_RAT | KVKRITPRHLQL | 77 | 88 | 4 |
| rChymotrypsin | H2AZ_RAT | QFPVGRIHRHL | 26 | 36 | 4 |

| | | | | | |
|---|---|---|---|---|---|
| rChymotrypsin | H2AZ_RAT | QLAIRGDEELDSL | 87 | 99 | 4 |
| rChymotrypsin | H2B1A_RAT | ERIAGEASRL | 72 | 81 | 4 |
| rChymotrypsin | H2B1A_RAT | ERIAGEASRLAHY | 72 | 84 | 4 |
| rChymotrypsin | H2B1A_RAT | E[-18.0106]RIAGEASRL | 72 | 81 | 4 |
| rChymotrypsin | H2B1A_RAT | E[-18.0106]RIAGEASRLAHY | 72 | 84 | 4 |
| rChymotrypsin | H2B1A_RAT | KQVHPDTGISSK[56.0262]AM[15.9949]S[27.9949]IM | 47 | 63 | 4 |
| rChymotrypsin | H2B1A_RAT | KQVHPDTGISS[-18.0106]K[70.0419]AM[15.9949]SIM[15.9949] | 47 | 63 | 3 |
| rChymotrypsin | H2B1A_RAT | KQVHPDTGIS[-18.0106]S[-18.0106]K[56.0262]AMS[-18.0106]IMNSF | 47 | 66 | 3 |
| rChymotrypsin | H2B1A_RAT | KVLKQVHPDTGISSK[270.1441]AMSIM[15.9949] | 44 | 63 | 2 |
| rChymotrypsin | H2B1A_RAT | KVLKQVHPDTGISSK[270.1441]AM[15.9949]SIM[15.9949] | 44 | 63 | 3 |
| rChymotrypsin | H2B1A_RAT | KVLKQVHPDTGISSK[56.0262]AM[15.9949]S[27.9949]IM[15.9949] | 44 | 63 | 4 |
| rChymotrypsin | H2B1A_RAT | KVLKQVHPDTGISS[27.9949]K[56.0262]AMSIM | 44 | 63 | 2 |
| rChymotrypsin | H2B1A_RAT | KVLKQVHPDTGISS[27.9949]K[56.0262]AM[15.9949]SIM | 44 | 63 | 3 |
| rChymotrypsin | H2B1A_RAT | KVLKQVHPDTGIS[-18.0106]S[-18.0106]K[42.0470]AM[15.9949]S[27.9949]IM[15.9949] | 44 | 63 | 3 |
| rChymotrypsin | H2B1A_RAT | PEVSAKGTTISKKGF | 1 | 15 | 4 |
| rChymotrypsin | H2B1_RAT | AHYNKRSTITSREIQTAVRL | 80 | 99 | 4 |
| rChymotrypsin | H2B1_RAT | AKHAVSEGTKAVTKY | 106 | 120 | 4 |
| rChymotrypsin | H2B1_RAT | GIMNSFVNDIF | 60 | 70 | 4 |
| rChymotrypsin | H2B1_RAT | GIM[15.9949]NSFVNDIF | 60 | 70 | 4 |
| rChymotrypsin | H2B1_RAT | KQVHPDTGISSKAM | 46 | 59 | 4 |
| rChymotrypsin | H2B1_RAT | KQVHPDTGISSKAMGIM | 46 | 62 | 4 |
| rChymotrypsin | H2B1_RAT | KQVHPDTGISSKAMGIMNSF | 46 | 65 | 4 |
| rChymotrypsin | H2B1_RAT | KQVHPDTGISSKAMGIMNS[-18.0106]F | 46 | 65 | 4 |
| rChymotrypsin | H2B1_RAT | KQVHPDTGISSKAMGIM[15.9949] | 46 | 62 | 4 |
| rChymotrypsin | H2B1_RAT | KQVHPDTGISSKAMGIM[15.9949]NSF | 46 | 65 | 4 |

| | | | | | |
|---|---|---|---|---|---|
| rChymotrypsin | H2B1_RAT | KQVHPDTGISSKAM[15.9949] | 46 | 59 | 4 |
| rChymotrypsin | H2B1_RAT | KQVHPDTGISSKAM[15.9949]GIM | 46 | 62 | 4 |
| rChymotrypsin | H2B1_RAT | KQVHPDTGISSKAM[15.9949]GIMNSF | 46 | 65 | 4 |
| rChymotrypsin | H2B1_RAT | KQVHPDTGISSKAM[15.9949]GIMNS[-18.0106]F | 46 | 65 | 4 |
| rChymotrypsin | H2B1_RAT | KQVHPDTGISSKAM[15.9949]GIM[15.9949] | 46 | 62 | 4 |
| rChymotrypsin | H2B1_RAT | KQVHPDTGISSKAM[15.9949]GIM[15.9949]NSF | 46 | 65 | 4 |
| rChymotrypsin | H2B1_RAT | KVLKQVHPDTGISSKAM | 43 | 59 | 4 |
| rChymotrypsin | H2B1_RAT | KVLKQVHPDTGISSKAMGIM | 43 | 62 | 4 |
| rChymotrypsin | H2B1_RAT | KVLKQVHPDTGISSKAMGIM[15.9949] | 43 | 62 | 4 |
| rChymotrypsin | H2B1_RAT | KVLKQVHPDTGISSKAM[15.9949] | 43 | 59 | 4 |
| rChymotrypsin | H2B1_RAT | KVLKQVHPDTGISSKAM[15.9949]GIM | 43 | 62 | 4 |
| rChymotrypsin | H2B1_RAT | KVLKQVHPDTGISSKAM[15.9949]GIM[15.9949] | 43 | 62 | 4 |
| rChymotrypsin | H2B1_RAT | KVLKQVHPDTGIS[-18.0106]SKAM | 43 | 59 | 4 |
| rChymotrypsin | H2B1_RAT | KVLKQVHPDTGIS[-18.0106]SKAMGIM | 43 | 62 | 4 |
| rChymotrypsin | H2B1_RAT | KVLKQVHPDTGIS[-18.0106]SKAM[15.9949] | 43 | 59 | 4 |
| rChymotrypsin | H2B1_RAT | KVLKQVHPDTGIS[-18.0106]SKAM[15.9949]GIM | 43 | 62 | 3 |
| rChymotrypsin | H2B1_RAT | KVLKQVHPDTGIS[-18.0106]S[-18.0106]KAM | 43 | 59 | 4 |
| rChymotrypsin | H2B1_RAT | K[114.0317]VLKQVHPDTGISSKAM[15.9949]GIM | 43 | 62 | 4 |
| rChymotrypsin | H2B1_RAT | K[27.9949]VLKQVHPDTGISSKAM | 43 | 59 | 3 |
| rChymotrypsin | H2B1_RAT | K[27.9949]VLKQVHPDTGISSKAM[15.9949] | 43 | 59 | 2 |
| rChymotrypsin | H2B1_RAT | LPGELAKHAVSEGTKAVTKY | 101 | 120 | 4 |
| rChymotrypsin | H2B1_RAT | LPGELAK[42.0470]HAVSEGTK[70.0419]AVTKY | 101 | 120 | 4 |
| rChymotrypsin | H2B1_RAT | NKRSTITSREIQTAVRL | 83 | 99 | 4 |
| rChymotrypsin | H2B1_RAT | NKRSTITSREIQTAVRLL | 83 | 100 | 4 |
| rChymotrypsin | H2B1_RAT | NKRS[-18.0106]TITSREIQTAVRL | 83 | 99 | 4 |
| rChymotrypsin | H2B1_RAT | NK[27.9949]RSTITSREIQTAVRL | 83 | 99 | 3 |
| rChymotrypsin | H2B1_RAT | PGELAKHAVSEGTKAVTKY | 102 | 120 | 4 |
| rChymotrypsin | H2B1_RAT | VYKVLKQVHPDTGISSKAM[15.9949] | 41 | 59 | 3 |
| rChymotrypsin | H31_RAT | ALQEACEAY | 91 | 99 | 4 |
| rChymotrypsin | H31_RAT | QSSAVM[15.9949]AL | 85 | 92 | 4 |
| rChymotrypsin | H31_RAT | QSSAVM[15.9949]ALQEACEAY | 85 | 99 | 4 |
| rChymotrypsin | H31_RAT | RFQSSAVM | 83 | 90 | 4 |

| | | | | | |
|---|---|---|---|---|---|
| rChymotrypsin | H33_RAT | EDTNLCAIHAKRVTIMPKDIQL | 105 | 126 | 4 |
| rChymotrypsin | H33_RAT | EDTNLCAIHAKRVTIM[15.9949]PKDIQL | 105 | 126 | 4 |
| rChymotrypsin | H33_RAT | E[-18.0106]DTNLCAIHAKRVTIMPKDIQL | 105 | 126 | 4 |
| rChymotrypsin | H33_RAT | E[-18.0106]DTNLCAIHAK[56.0262]R[10.0083]VTIMPKDIQL | 105 | 126 | 3 |
| rChymotrypsin | H33_RAT | IRKLPF | 62 | 67 | 4 |
| rChymotrypsin | H33_RAT | IRKLPFQRL | 62 | 70 | 4 |
| rChymotrypsin | H33_RAT | KTDLRF | 79 | 84 | 4 |
| rChymotrypsin | H33_RAT | K[14.0156]TDLRF | 79 | 84 | 4 |
| rChymotrypsin | H33_RAT | K[28.0313]TDLRF | 79 | 84 | 4 |
| rChymotrypsin | H33_RAT | LIRKLPF | 61 | 67 | 4 |
| rChymotrypsin | H33_RAT | QKSTELL | 55 | 61 | 4 |
| rChymotrypsin | H33_RAT | QKS[-18.0106]TELL | 55 | 61 | 4 |
| rChymotrypsin | H33_RAT | QRLVREIAQDF | 68 | 78 | 4 |
| rChymotrypsin | H33_RAT | QRLVREIAQDFKTDL | 68 | 82 | 4 |
| rChymotrypsin | H33_RAT | QRLVREIAQDFK[28.0313]TDL | 68 | 82 | 4 |
| rChymotrypsin | H33_RAT | QSAAIGALQEASEAY | 85 | 99 | 4 |
| rChymotrypsin | H33_RAT | QSAAIGALQEAS[-18.0106]EAY | 85 | 99 | 4 |
| rChymotrypsin | H33_RAT | QS[-18.0106]AAIGALQEASEAY | 85 | 99 | 4 |
| rChymotrypsin | H33_RAT | RFQSAAIGALQEASEAY | 83 | 99 | 4 |
| rChymotrypsin | H33_RAT | RPGTVALREIRRY | 42 | 54 | 4 |
| rChymotrypsin | H33_RAT | VREIAQDF | 71 | 78 | 4 |
| rChymotrypsin | H33_RAT | VREIAQDFKTDL | 71 | 82 | 4 |
| rChymotrypsin | H33_RAT | VREIAQDFKTDLRF | 71 | 84 | 4 |
| rChymotrypsin | H33_RAT | VREIAQDFK[14.0156]TDL | 71 | 82 | 4 |
| rChymotrypsin | H33_RAT | VREIAQDFK[14.0156]TDLRF | 71 | 84 | 4 |
| rChymotrypsin | H33_RAT | VREIAQDFK[28.0313]TDL | 71 | 82 | 3 |
| rChymotrypsin | H33_RAT | VREIAQDFK[28.0313]TDLRF | 71 | 84 | 4 |
| rChymotrypsin | H4_RAT | ALKRQGRTLY | 89 | 98 | 4 |
| rChymotrypsin | H4_RAT | ARRGGVKRISGL | 38 | 49 | 4 |
| rChymotrypsin | H4_RAT | ARRGGVKRISGLIY | 38 | 51 | 4 |
| rChymotrypsin | H4_RAT | EETRGVL | 52 | 58 | 4 |
| rChymotrypsin | H4_RAT | EETRGVLKVF | 52 | 61 | 4 |

| | | | | | |
|---|---|---|---|---|---|
| rChymotrypsin | H4_RAT | ENVIRDAVTY | 63 | 72 | 4 |
| rChymotrypsin | H4_RAT | E[-18.0106]ETRGVL | 52 | 58 | 3 |
| rChymotrypsin | H4_RAT | E[-18.0106]NVIRDAVTY | 63 | 72 | 4 |
| rChymotrypsin | H4_RAT | IYEETRGVL | 50 | 58 | 4 |
| rChymotrypsin | H4_RAT | IYEETRGVLKVF | 50 | 61 | 4 |
| rChymotrypsin | H4_RAT | KRQGRTLY | 91 | 98 | 4 |
| rChymotrypsin | H4_RAT | KVFLENVIRDAVTY | 59 | 72 | 4 |
| rChymotrypsin | H4_RAT | LENVIRDAVTY | 62 | 72 | 4 |
| rChymotrypsin | H4_RAT | RDNIQGITKPAIRRLARRGGVKRISGL | 23 | 49 | 4 |
| rChymotrypsin | H4_RAT | TEHAKRKTVTAMDVVY | 73 | 88 | 4 |
| rChymotrypsin | H4_RAT | TEHAKRKTVTAM[15.9949]DVVY | 73 | 88 | 4 |

**Table S5:** Per-Sample Reagent and Researcher Time Cost Comparison (5 µg Histone Peptide Input)*

| Workflow | Reagent/Sample | Prep Time/Batch | Researcher/Sample | MS Runs/Sample | Total/Sample | Total/100 Samples |
|---|---|---|---|---|---|---|
| Trypsin + propionylation | $0.70 | 6–12 h | $24.00–$48.00 | 1 | **$24.70–$48.70** | **$2,470–$4,870** |
| Arg-C Ultra + propionylation | $1.34 | ~6 h | $24.00 | 1 | **$25.34** | **$2,534** |
| Arg-C Ultra (unlabeled) | $1.33 | ~2 h | $8.00 | 1 | **$9.33** | **$933** |
| Arg-C Ultra + TMTzero | $4.58 | ~3 h | $12.00 | 1 | **$16.58** | **$1,658** |
| Arg-C Ultra + TMT10plex | $131.50 | ~3 h | $12.00 | 0.1 | **$143.50** | **$14,350** |
| RIPUP + propionylation | $1.34+ | ~6 h | $24.00 | 2 | **$25.34+** | **$2,534+** |

| | | | | | | |
|---|---|---|---|---|---|---|
| RIPUP (unlabeled) | $1.33+ | ~2 h | $8.00 | 2 | **$9.33+** | **$933+** |
| RIPUP + TMTzero | $4.58+ | ~3 h | $12.00 | 2 | **$16.58+** | **$1,658+** |
| RIPUP + TMT10plex | $131.50+ | ~3 h | $12.00 | 0.2 | **$143.50+** | **$14,350+** |

* All costs are reported in USD and exclude instrument acquisition time; the MS Runs/Sample column is provided so that readers may calculate instrument costs using local rates. Reagent costs were calculated from manufacturer list prices assuming 5 µg histone peptide input per sample. Enzyme prices: Trypsin Gold MS-grade (Promega, $138/100 µg) at a 1:10 enzyme-to-substrate (E:S) ratio ($0.69/sample); Arg-C Ultra (Promega, $133/5 µg) at a 1:100 E:S ratio ($1.33/sample). r-Chymotrypsin (Promega, Early Access program) pricing has not been established; RIPUP workflow totals are reported as additive to the Arg-C Ultra cost. TMTzero (Thermo Scientific, Cat. No. 90067, $325/5 x 0.8 mg) is costed for single-channel derivatization, with per-sample cost assuming proportional scaling of the manufacturer-recommended protocol (0.8 mg per 100 µg peptide) to 5 µg input (~20 samples per vial, ~100 samples per kit, $3.25/sample). TMT10plex (Thermo Scientific, Cat. No. 90110, $1,302/10 × 0.8 mg) is costed for standard 10-plex isobaric multiplexing ($130.17/sample); ten samples are combined per MS run. Propionic anhydride (Sigma-Aldrich, Cat. No. 240311, ≥99%, ~$30–50/100 mL) per-sample reagent cost is negligible (<$0.01). Researcher time was estimated at $40/h for batches of 10 samples processed in parallel, with per-sample cost calculated as (batch prep time ÷ 10) x $40. Propionylation workflows include two pre-digestion and two post-digestion derivatization rounds and enzymatic digestion; the trypsin-based workflow spans two calendar days due to overnight digestion (6–12 h range reflects variability in digestion time and batch complexity). Arg-C Ultra and RIPUP workflows without chemical derivatization (~2 h) include digestion; RIPUP runs Arg-C Ultra and r-Chymotrypsin digestions in parallel on separate aliquots. TMT labeling adds ~1 h (reagent reconstitution, incubation, hydroxylamine quench). Non-multiplexed workflows require one MS run per sample per protease (1 for single protease, 2 for RIPUP dual protease). TMT10plex workflows combine 10 samples per run (0.1 for single protease, 0.2 for RIPUP). All reagent prices reflect manufacturer list prices as of mid-2026 and may vary with institutional contracts or volume discounts. Costs do not include instrument acquisition time, consumables (e.g., desalting tips, LC columns), or sample extraction.
